# Differential expression of GABARAPs in GBM renders temozolomide sensitivity in a p53-dependent manner

**DOI:** 10.1101/2024.10.17.618809

**Authors:** Megha Chaudhary, Naveen Soni, Shweta Dongre, Ashwani Tiwari, Nargis Malik, Kunzang Chosdol, Bhawana Bissa

## Abstract

Glioblastoma (GBM) is one of the most debilitating and extremely aggressive tumors, with a median survival of less than a year. GBMs have high metastatic potential and frequently acquire chemoresistance. The current multimodal treatment approaches for GBM include surgical tumor resurrection, radiotherapy, and chemotherapy but these approaches leave the patient with long-term disabilities such as depletion of cognitive abilities, leukoencephalopathy, and recurrence in 6-8 months. Glioma cells are highly dependent on autophagy to survive and proliferate. Autophagy inhibition has proven to be a beneficial strategy for restricting glioma growth. However, the autophagy pathway cannot be efficiently targeted due to the lack of specific autophagy inhibitors. Understanding the vulnerabilities in autophagy gene expression can help to design better autophagy inhibitors. This study demonstrates the differential expression of GABARAP family members in low-grade glioma and GBM. Our study highlights the differential expression of GABARAP family members in response to autophagy inhibition and induction. Moreover, the knockdown of specific GABARAP family members enhanced proliferation and reduced temozolomide (TMZ) sensitivity of glial cells by decreasing the p53 expression. The selective expression pattern of GABARAP genes in Glioblastoma can be utilized to screen for patients who might respond better to temozolomide treatment. The differential expression of GABARAP family members highlights the subtle regulation of the autophagy pathway in response to environmental cues.

## Introduction

Autophagy is an evolutionarily conserved intracellular process that occurs ubiquitously in all the eukaryotic cells^1^. It occurs at a basal level to clear damaged organelles and turnover of proteins to maintain cellular homeostasis. Some studies connect autophagy with the life span extension, but due to its dual role in disease progression, it has been difficult to solve this paradox^2^. Autophagy has been implicated in the vast web of diseases such as neurodegenerative disorders, non-alcoholic fatty liver disease (NAFLD), inflammatory bowel disease (IBD), acute kidney disease (AKD), chronic kidney disease (CKD), diabetes, and cancer^3^. It has been demonstrated that autophagy regulates tumors in two ways. It suppresses carcinogenesis and limits the ability of cancer cells to proliferate, but it also encourages the growth of tumor cells^4,5^. The autophagy pathway involves ∼50 ATG proteins that were discovered individually by genetic screening of their mutants. Some of these are the ATG8 family of proteins that are concerned with the recruitment of cargo, autophagosome biogenesis, its transport, and fusion with the lysosome^6^. The ATG8 family has 8 orthologs; LC3A, LC3B, LC3B2, LC3C (encoded by Microtubule-Associated Protein-1 Light Chain-3 or MAP1LC3), and γ-aminobutyric acid receptor-associated protein (GABARAP), GABARAPL1 (GABARAP like protein 1), GABARAPL2 (also known as GATE-16), and GABARAPL3^7,8^. The role of ATG8 proteins has been studied in yeast, where they participate in membrane trafficking and autophagy^8^. Additionally, non-autophagic functions of ATG8 and their involvement in aging, neurodegenerative disorders, and in the cell secretome (extracellular vesicles) have also been discovered^9–12^. Moreover, some previous studies have found a good prognostic role of GABARAPs ^13–19^ and MAP1LC3 genes^13,20–24^.

Glioblastoma (GBM) is the most common, aggressive, and malignant astrocytoma, which occurs in cerebral hemispheres, mostly in the supratentorial region of CNS^25^. It appears as either grade-IV astrocytoma (90%) or progression of lower-grade astrocytoma (10%)^26^. On an average the median survival time is just 12 to 18 months, with 25% of patients surviving longer than a year and 5% surviving longer than five years after initial diagnosis^27^. The primary reasons for GBM are thought to be some genetic alterations that cause aberrant metabolism in progenitor or neuroglial cells. Despite the wide range of current therapies, there is a challenge in GBM treatment due to 1) Tumor-induced seizures 2) Inborn resistance to conventional therapies 3) High multidrug resistance (MDR) arising by cancer stem cells and 4) High migration to nearby tissues^28,29^. The standard care for GBM is surgical resection followed by a combined treatment of radiotherapy and alkylating agent temozolomide for multiple cycles^30^.

Temozolomide (TMZ) is a DNA alkylating agent that methylates DNA specifically at the O^6^ position of guanines, causing cell cycle arrest at the G2/M checkpoint^31,32^. It is an oral drug approved by the FDA as a first line of defense in grade III and grade IV glioma^33^. Several studies determined an overall 2.5 months of increased median survival with TMZ treatment, but due to high activity of O^6^-Methylguanine-DNA methyl transferase (MGMT), TMZ resistance has become a major clinical challenge^34^. In addition, TMZ resistance in MGMT-deficient GBMs is one of the conflicting outcomes that mark the presence of another component to induce TMZ resistance^34^.

Chloroquine (CQ) is a well-known antimalarial medication, although its current use is diminished due to the advent of CQ-resistant strains of the malarial parasite ^35^. The early spark of CQ’s anti-neoplastic property was detected in the 1970s when the incidence of Burkitt’s lymphoma was dramatically reduced in anti-malarial experiments^36^. After a series of observations, it was reported that CQ has the potential to sensitize cells against radiation and chemotherapy in a wide spectrum of cancers, including GBM^37^. At the physiological pH of 7.4, an unprotonated form of CQ efficiently penetrates the cell, gets reserved in acidic compartments, and becomes protonated. Consequently, it raises intra compartmental pH and disrupts normal endosome, autophagosome, and lysosome fusion activities leading to disruption of autophagy^38^. Previous phase I/II clinical trials used CQ along with conventional therapy (resection, radiation, and TMZ treatment) and have shown significantly increased median survival time up to 25 months^39–41^. Low-grade glioma (LGG) is the primary brain tumor that is an early-grade brain tumor, with a higher survival rate compared to GBM^42^. In this study, we analyzed the differential expression of

GABARAP family members in GBM and LGG patient samples. Further, we investigated how CQ alters GABARAP expression and enhances TMZ-mediated cytotoxicity in glioma cell lines. We identified that GABARAP mRNA and protein expression was significantly increased upon CQ treatment. Temozolomide treatment differentially enhanced GABARAP mRNA and protein expression, while reducing the expression of MAP1LC3A. Previous study demonstrates that TP53 acts as rheostat and helps maintain autophagic homeostasis^43^. TP53 was found to have a positive correlation with GABARAP and we observed that GABARAP was essential in maintaining steady-state levels of TP53 under TMZ treatment. Importantly, we also demonstrated that knockdown and overexpression of GABARAP regulated TMZ sensitivity in glial cell line, and combined treatment of CQ and TMZ increased the expression of TP53 suppressing the proliferation of glial cell lines. Based on our study we speculate that GBM patients having high GABARAP expression would probably respond better to temozolomide treatment and consequently have better overall survival.

## Results

### 1. ATG8 genes show varied expression levels in GBM and LGG patient database

ATG8 genes are associated with the autophagy pathway as well as in an array of non-autophagic pathways, including cell trafficking, modulation of cellular transport, endocytosis, cell signaling and extracellular vesicle biogenesis and secretion^44,45^. By using the GEPIA^46^ data set, we examined the expression of all ATG8 orthologs (MAP1LC3A, MAP1LC3B, MAP1LC3C, GABARAP, GABARAPL1, GABARAPL2) in GBM and LGG patient samples (Fig. 1). GBM and LGG are two distinct late and early-stage glioma cancer stages respectively. We examined how ATG8 genes were expressed in these two different stages. In GBM and LGG, GABARAP was significantly upregulated (Fig 1A), while GABARAPL1 was significantly downregulated (Fig 1B). However, GABARAPL2 did not show a significant difference. Among MAP1LC3 genes, MAP1LC3A was significantly downregulated in GBM and LGG samples (Fig 1D). While MAP1LC3B did not show a significant difference, MAP1LC3C showed a significant upregulation in GBM but not in LGG samples (Fig 1E and 1F). Further, by using Correlation AnalyzeR^47^, we analyzed the co-expression of these genes and generated a principal component analysis (PCR) plot of ATG8 genes in normal and brain cancer (Fig 1G, 1H). PCA plot principal component-1 (PC1) and principal component-2 (PC2) accounted for 70 to 80% of the entire variation in gene expression. In both normal and tumor conditions, scattering patterns of all ATG8 genes were different. During normal conditions, GABARAPL1 and MAP1LC3A were grouped together and GABARAP, GABARAPL2, and MAP1LC3B were in close proximity (Fig 1G) and MAP1LC3C gene did not form any cluster. However, in tumor conditions, ATG8 family members behave differently and do not form any cluster, which highlights the variation in functionality of different members during glioma progression^48^. Since GBM tumors are classified into four subclasses: Proneural (P), Neural (N), Classical (C), and Mesenchymal (M), ATG8 genes were regulated according to these subclasses (Supp. Fig 1A to 1C and 1G to 1I). We used multivariate Kaplan–Meier survival analysis plots to assess the overall survival effect of ATG8 gene expression. We observed that while GABARAP was positively correlated with overall survival (Supp. Fig 1D), MAP1LC3A was negatively correlated with overall survival (Supp Fig 1J), highlighting their differential effect on GBM survival. These results highlight the unique role that each ATG8 gene might play in the progression of glioma therefore we further wanted to confirm their expression in Indian Glioma patient samples collected from AIIMS, New Delhi.

**Figure 1.**
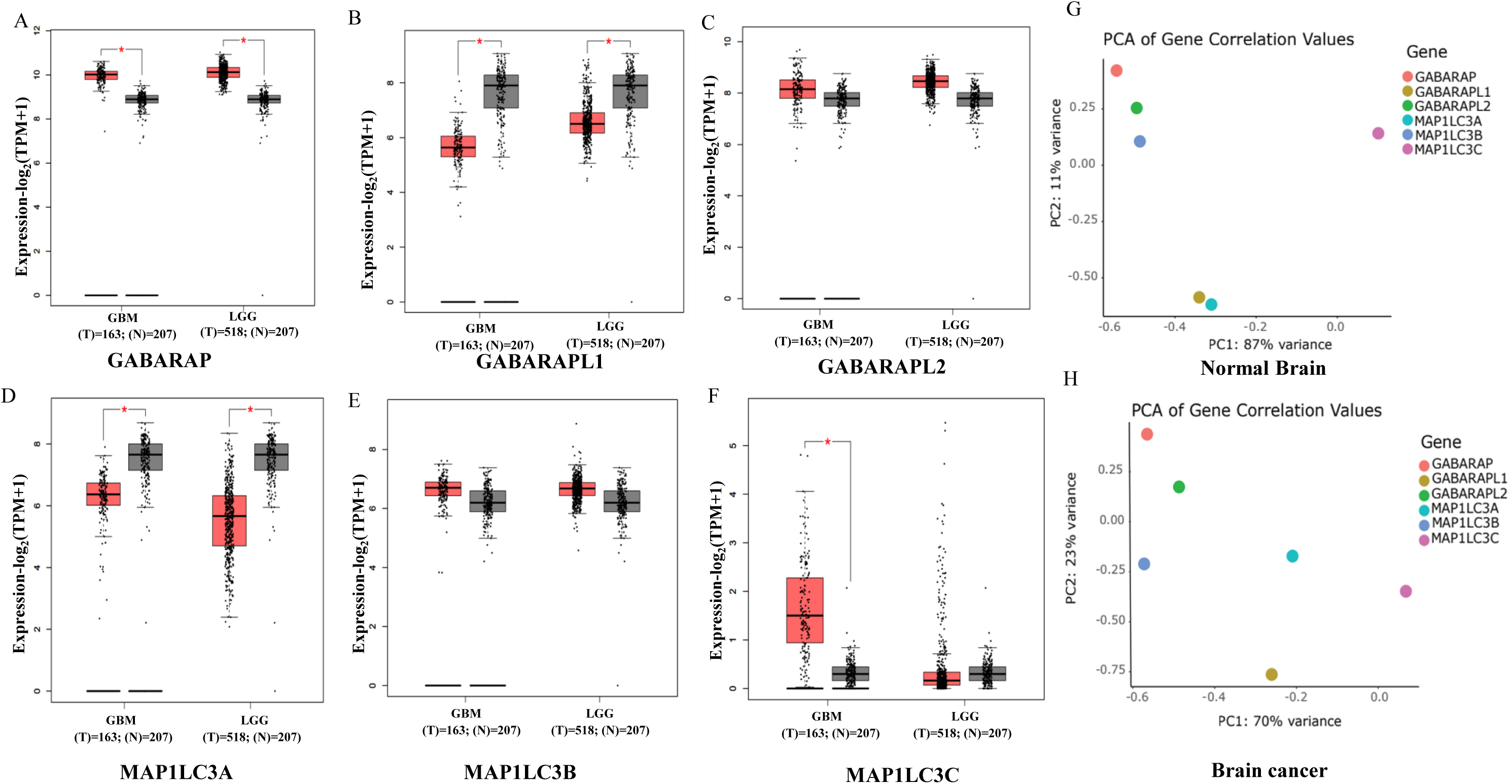
Expression of ATG8 in normal and glioma (A to F) Expression box plot for GABARAP’s (GABARAP, GABARAPL1, GABARAPL2), and LC3 subfamily (MAP1LC3A, MAP1LC3B, and MAP1LC3C) show differential expression in GBM and LGG by using GEPIA analysis. log2(TPM+1) is the log2 of the Transcript Count Per Million. (G and H) Correlation analysis of the expression of GABARAPs and MAP1LC3s in glioma and normal brain by using Principal component analysis (PCA). Normal brain shows 11% variance while brain cancer showed 23% variance in ATG8.

### 2. ATG8 genes show different expression levels in Indian Glioma patient samples

To validate the patterns of ATG8 genes from in-silico RNA sequencing data, we analyzed their expression in GBM and LGG patient samples collected from AIIMS, New Delhi, after ethical clearance. There was no discrimination based on the ethnicity, gender, and age of the patients. In total, RNA samples from 10 GBM patients (5 male and 5 female) with a mean age of 37 years and 9 LGG patients (5 male and 4 female) with a mean age of 38.11 years were used for analysis. Normal brain RNA was used as a control sample to compare expression levels. A summary of all subjects that are studied in this work is provided in Table. Firstly, we analyzed the mRNA expression of all ATG8 genes in GBM and LGG patient samples and compared them with normal brain RNA. We performed real-time PCR and observed that expression of the GABARAP gene was significantly elevated in 7/10 GBM samples (Fig 2A and Supp Fig 2A) as previously observed in in-silico data (Fig 1A). However, GABARAP was not significantly altered in LGG patient samples. Further, we observed significant downregulation of GABARAPL1 (8/10) in GBM samples but not in LGG samples (Fig 2B and Supp Fig 2A and 2B) GABARAPL2 did not show significant alteration in GBM and LGG patient samples (Fig 2C). On further analysis we observed an increased but non-significant change in the expression of MAP1LC3A in GBM and LGG patient samples (Fig 2D), which contrasted concerning the in-silico data, observed in Fig 1D. Contrary to the in-silico data in Fig 1E, we observed a significant reduction in MAP1LC3B levels in GBM and LGG patient samples (Fig 2E). Finally, we analyzed MAP1LC3C levels and observed wide variation in its expression in GBM and LGG patient samples (Fig 1F), which might be due to the expression of the splice variant (280 bp instead of 154 bp) as observed in gel pictures (Supp Fig2A and 2B). To verify the expression of individual ATG8 genes we checked their expression in Glial cell lines U87MG and LN229 as well as in normal brain samples and as expected we observed the expression of all ATG8 genes except MAP1LC3C (Supp Fig 2C). Thus, we observed wide variation in the expression of ATG8 genes in the Glioma patient database as well as Indian glioma patient samples which highlights that different ATG8s might play unique roles in glioma progression, autophagy and chemosensitivity.

**Figure 2.**
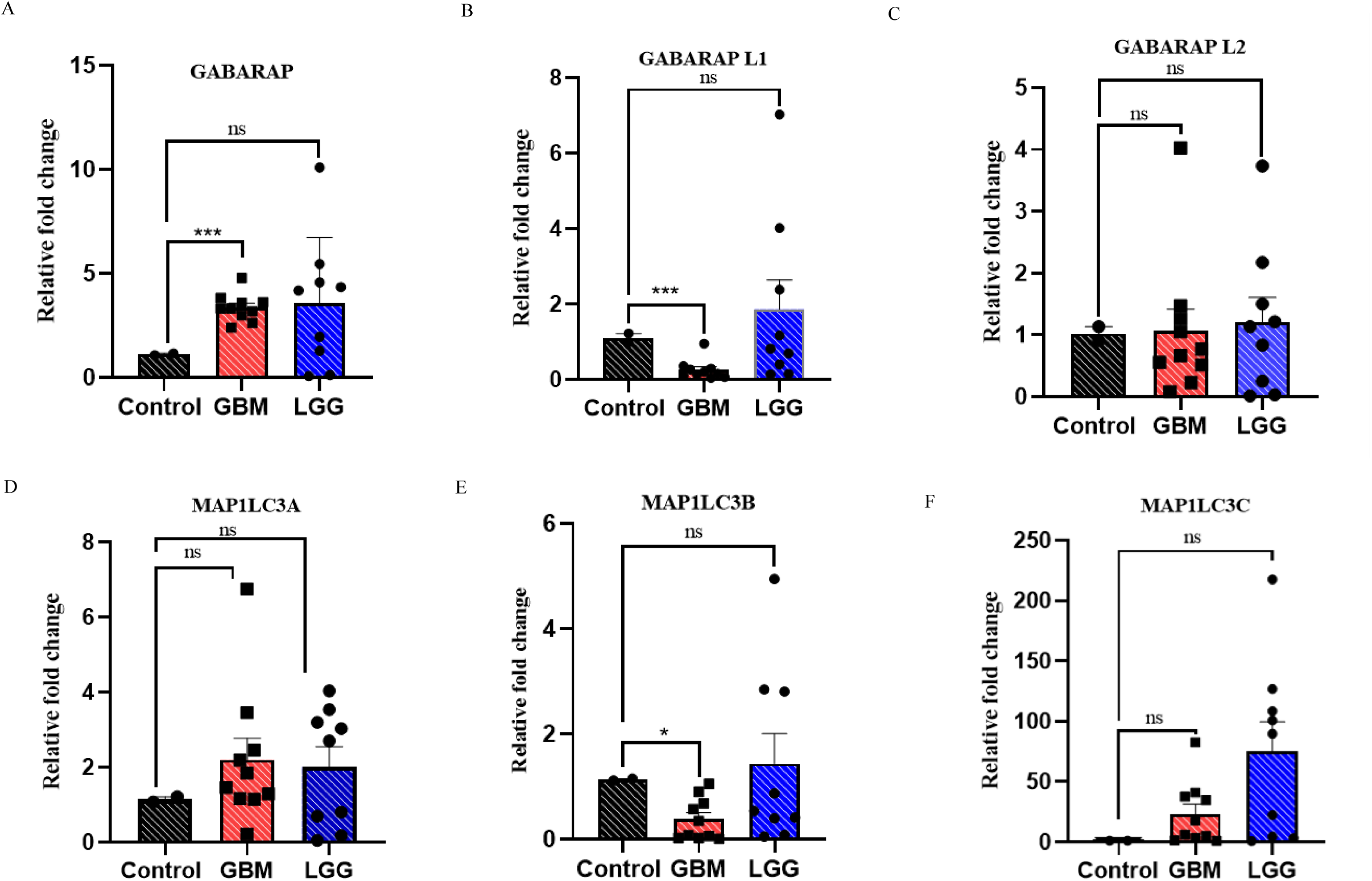
Expression analysis of GABARAP and MAP1LC3 family members in GBM and LGG Patient samples. Quantification of GABARAP (A-C) and MAP1LC3 (D-F) family members using qRT-PCR in GBM and LGG patient samples compared to normal brain samples. GABARAP gene expression is significantly increased while GABARAPL1 and MAP1LC3B expression is significantly reduced in GBM patient samples. All the statistical values were quantified using an unpaired student t-test (* p< 0.05; **p <0.01, ***p <0.001).

### 3. GABARAPs demonstrate different expression and lipidation in response to autophagy perturbation

Since GABARAPs showed significant differences in expression in glioma patient samples, we wanted to analyze their expression and lipidation in response to autophagy perturbation. To evaluate the effect of autophagy perturbation on GABARAP expression, we subjected glial cell lines to autophagy inhibition and induction. In vitro, several lysosomal inhibitors, including chloroquine (CQ), protease inhibitor (Pepstatin A and Leupeptin), and Bafilomycin A1 (BafA1), have been employed interchangeably to block autophagy flux^49^. CQ and its derivative hydroxychloroquine (HCQ) are FDA-approved medications and have recently been used in a clinical trial to improve cancer treatment by inhibiting autophagy^50,51^. We treated glial cell lines with a series of CQ concentrations and determined IC50 values of 36 μM and 43.45 μM in U87MG and LN229, respectively (Supp Fig. 3A). To evaluate the effect of autophagy perturbation on GABARAP’s mRNA expression, we treated U87MG and LN229 cells with 20μM CQ for 4hours. Real-time PCR showed a significant increase in mRNA expression of GABARAP by >1.5 fold in both U87MG and LN229 cells, however, no significant change in expression of GABARAPL1 and GABARAPL2 was observed (Fig. 3A, Supp Fig. 3B). To validate the expression levels, we ran the PCR product on gel and observed a significant increase in GABARAP transcript levels after CQ treatment (Fig 3A). We further analyzed the effect of autophagy induction by starvation on mRNA expression of GABARAP’s by incubating cells in earle’s balanced salt solution (EBSS), a widely used media for mimicking starvation. Interestingly, after 1 hour of EBSS treatment, the GABARAPL2 transcript level was significantly elevated compared to GABARAP and GABARAPL1 in both U87MG and LN229 (Fig 3B and Supp Fig 3C). The increase in GABARAP mRNA expression was also observed after treatment with BafilomycinA1 (Supp Fig 3D). To validate the transcript level changes at the protein level, we did a western blot to check the expression of the GABARAPs after treatment with 20μM CQ for 4 hours. Intriguingly, level of non-lipidated GABARAP (GABARAP-I) was significantly elevated by >4 fold after CQ treatment while there was little or no change in levels of non-lipidated forms of GARABAPL1 and GABARAPL2. However, we observed an increase in lipidated forms of GABARAPL1 and GABARAPL2 upon CQ treatment (Fig 3C). On contrary, upon EBSS treatment we observed a slight decrease in GABARAP and GABARAPL1 total levels, which was not significant, and a significant increase in non-lipidated form of GABARAPL2 (Fig. 3D). To understand the changes in the protein level of GABARAPs we quantified the changes in their non-lipidated forms and observed that GABARAP-I levels were increased upon CQ treatment while GABARAPL2-I levels were increased upon EBSS treatment, signifying the differential response of GABARAPs if autophagy flux is blocked or induced.

**Figure 3.**
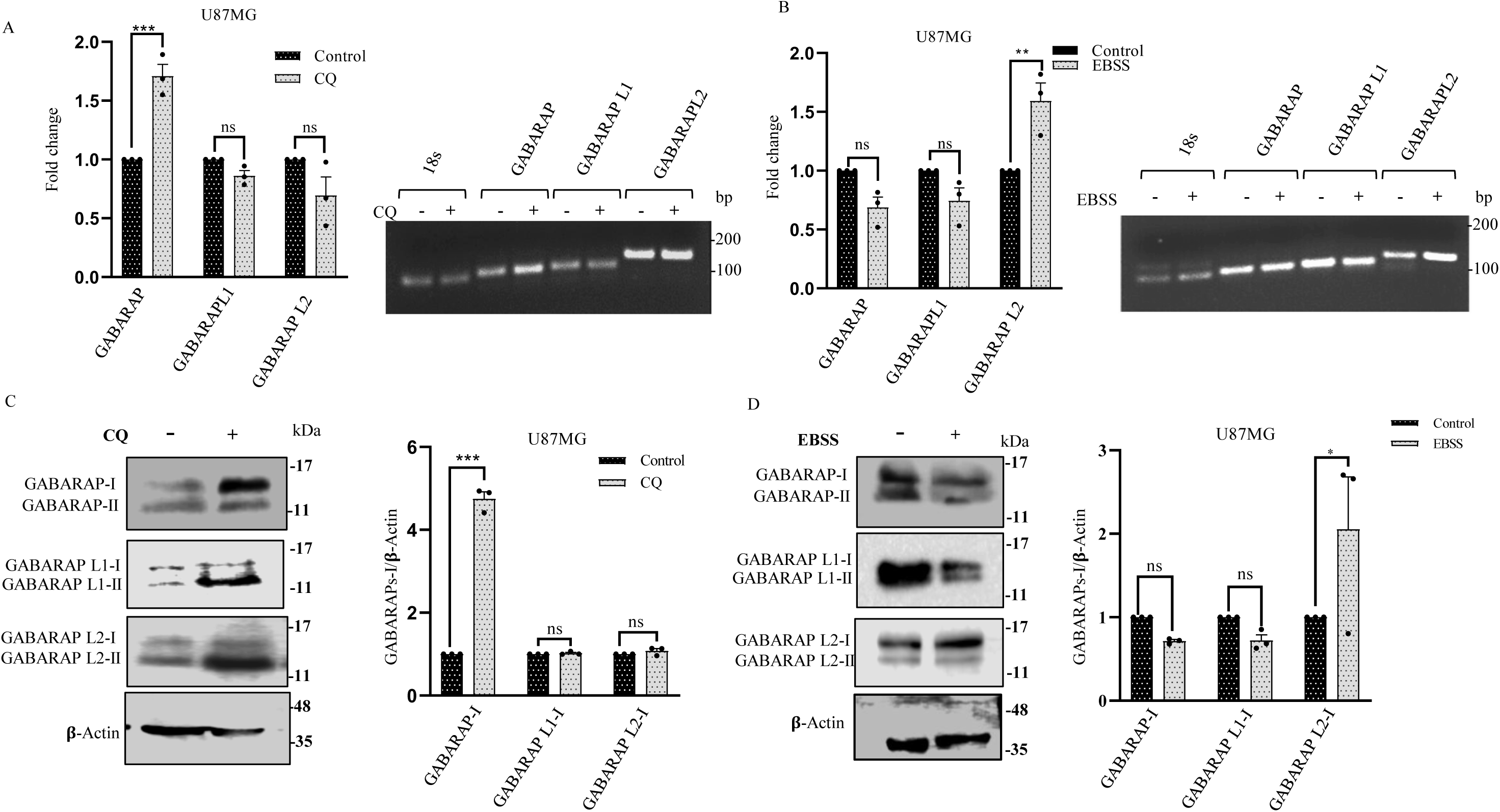
Differential expression of GABARAP family members upon autophagy modulation (A) Expression analysis of GABARAP, GABARAPL1, and GABARAPL2 after CQ drug treatment (20µM, 4 hours). A gel picture showing the increase in GABARAP expression after CQ treatment. (B) Expression analysis of GABARAP, GABARAPL1, and GABARAPL2 in U87MG cells after treatment with EBSS for 1 hour. A gel picture showing the increase in GABARAPL2 expression after EBSS treatment. Western blots of GABARAP, GABARAPL1, GABARAPL2 in U87MG cells treated with 20µM CQ for 4 hours (C) and EBSS for 1hour (D). β-actin was used as loading control. All the statistical values have been calculated using one-way ANOVA (* p< 0.05; **p <0.01, ***p <0.001).

### 4. Temozolomide enhances GABARAP expression in glioma cell line

Recent studies investigating the role of TMZ have concluded that it could induce autophagy in GBM and ultimately provide tumor plasticity and therapeutic resistance to tumor^52^. A previous study has reported that TMZ treatment enhances autophagy, and its inhibition greatly improves TMZ sensitivity. This study indicated that inhibition of autophagy may accelerate the outcome of TMZ-based cancer therapy^53^. However, the involvement of ATG8 proteins in TMZ sensitivity in GBM is not elucidated so in continuation of our study, we assessed how TMZ affects the expression of GABARAP and MAP1LC3 family. To check the TMZ cytotoxicity, we performed the MTT assay (Supp. Fig. 4A) and determined the IC50 for TMZ as 358.0 μM and 384.30 μM in U87MG and LN229, respectively. We treated U87MG with 358.0 μM TMZ and analyzed the expression of GABARAP and MAP1LC3A. A significant increase in GABARAP (Fig. 4A) and a significant decrease in MAP1LC3A expression (Fig. 4B) was observed. The expression of GABARAPL1, GABARAPL2, and MAP1LC3B was not altered significantly after TMZ treatment. Since TMZ alters G2/M checkpoint and arrest cells, it was noticed earlier that treatment with TMZ could enhance MAP1LC3B in a dose-dependent manner^54^. We were interested to know if the expression of GABARAP and MAP1LC3A genes was affected by TMZ in concentration dependent manner. After treatment of LN229 cells with increasing concentrations of TMZ, we observed an increase in GABARAP mRNA expression and a decrease in MAP1LC3A mRNA expression (Supp. Fig. 4B). Further, western blot analysis validated the time and concentration dependent increase in non-lipidated form of GABARAP levels after TMZ treatment (Fig 4C, 4D). Concomitantly, the expression of MAP1LC3A decreased with time and concentration-dependent treatment of TMZ. The reciprocal effect of TMZ on GABARAP and MAP1LC3A genes substantiates the effect of these genes on GBM patient survival as well (Supp Fig 1D, 1J). We also validated the effect of TMZ on autophagy by analyzing the expression of p62 and LC3B after TMZ treatment in LN229 cells. TMZ treatment caused decreased expression of p62, while there was an increase in the lipidated form of LCB (LC3B-II) (Supp. Fig. 4C), indicating autophagy induction after TMZ treatment. We further validated it by analyzing autophagy flux using tandem fusion of the red, acid-insensitive mCherry and the acid sensitive green fluorescent GFP double-tagged LC3B plasmid in LN229 cells. Confocal fluorescence microscopy demonstrated increased autophagy flux after TMZ treatment, as represented by an increased ratio of Red/Yellow dots (Supp Fig 4D). Thus, TMZ treatment increased autophagy flux and led to a differential effect on expression of GABARAP and MAP1LC3A.

**Figure 4:**
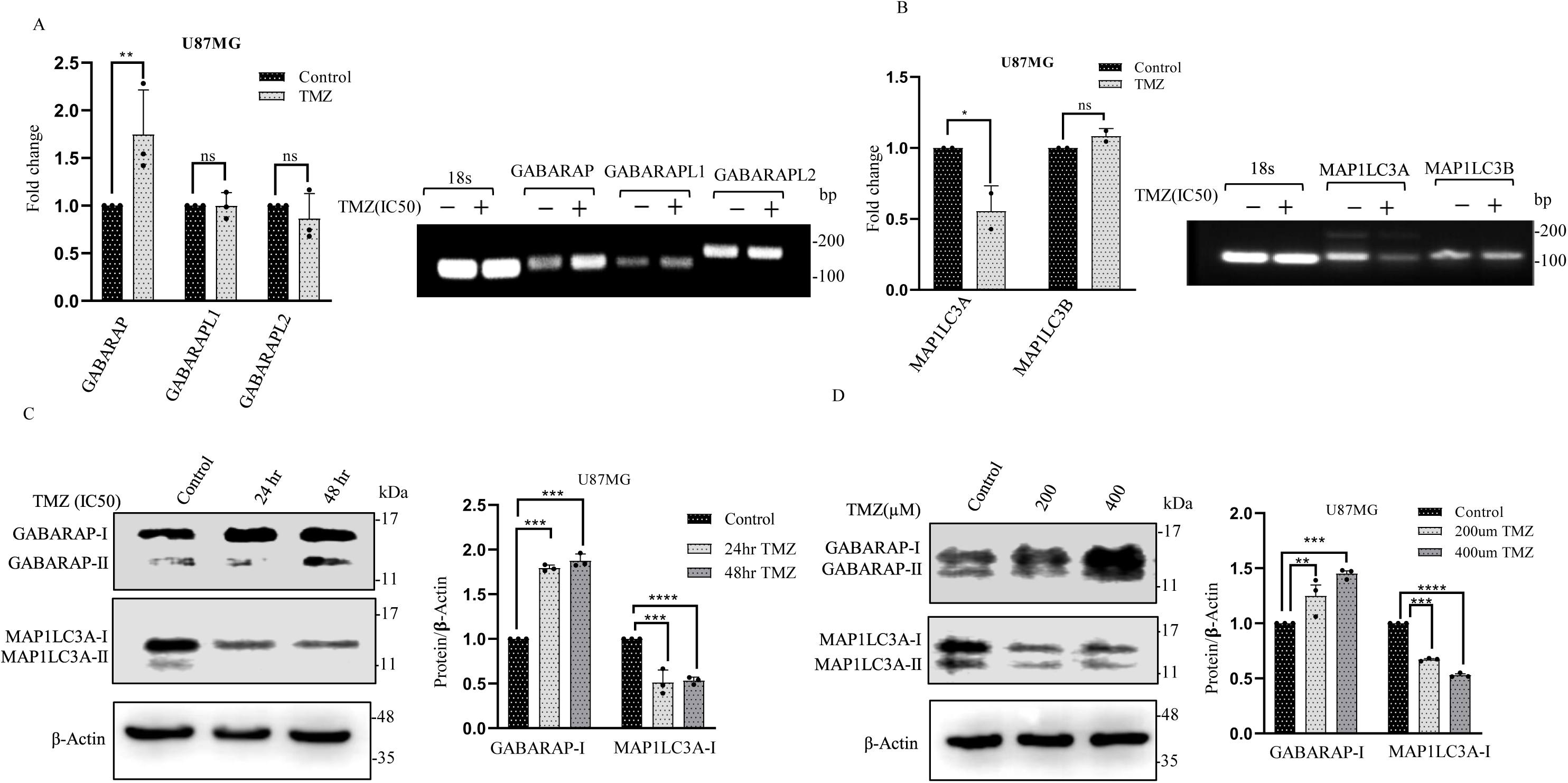
Expression analysis of GABARAP and MAP1LC3A after TMZ treatment (A-B) U87MG cells were treated with IC50 concentration of TMZ and analyzed for the expression of GABARAP, GABARAPL1, GABARAPL2, MAP1LC3A, and MAP1LC3B. (C-D) Western blot of GABARAP and MAP1LC3A in U87MG cell line treated with (C) IC50 TMZ concentration for 24hours and 48hours and (D) different concentrations of TMZ (200µM and 400µM) for 24 hours. β-actin used as loading control. All the statistical values have been calculated using one-way ANOVA (* p< 0.05; **p <0.01, ***p <0.001).

### 5. Knockdown of GABARAP enhances TMZ IC50 and suppresses p53 expression

To further investigate the mechanism of GABARAP in TMZ cytotoxicity we knocked down the expression of GABARAP using siRNA. In U87MG and LN229 cells, a significant knockdown of GABARAP was achieved compared to the control cells (Supp Fig. 5A). The same was validated at the protein level as well (Supp Fig. 5B). Knockdown of GABARAP increased the IC50 of TMZ in U87MG cells from 358.0μM to 496.40 μM (Fig 5A). Intriguingly, the increase in TMZ IC50 was specific to the knockdown of GABARAP and not GABARAPL1 and GABARAPL2 (Fig 5B). Moreover, the knockdown of GABARAP led to suppression of p53 expression in response to TMZ treatment, which was not observed with the knockdown of GABARAPL1 and GABARAPL2 (Fig 5C and Supp Fig 5C). We further validated the effect of GABARAP knockdown on p53 expression at the protein level and observed a significant decrease in p53 expression after TMZ treatment in GABARAP knockdown cells (Fig 5D). To examine whether GABARAP knockdown also affects autophagy flux, we treated control and GABARAP knockdown cells with CQ and, observed reduced accumulation of MAP1LC3B-II (Fig 5E) highlighting the important role of GABARAP in the robust flow of autophagy flux^39^. Finally, to investigate whether GABARAP siRNA affects the migration of U87MG, we performed a migration assay and observed that GABARAP knockdown with a combination of TMZ (IC50) does not significantly affect the cell migration (Supp. Fig. 5D). Thus, we could conclude that GABARAP might be essential in enhancing the cytotoxic effects of TMZ, probably by synergistically sustaining p53 levels upon TMZ treatment.

**Figure 5.**
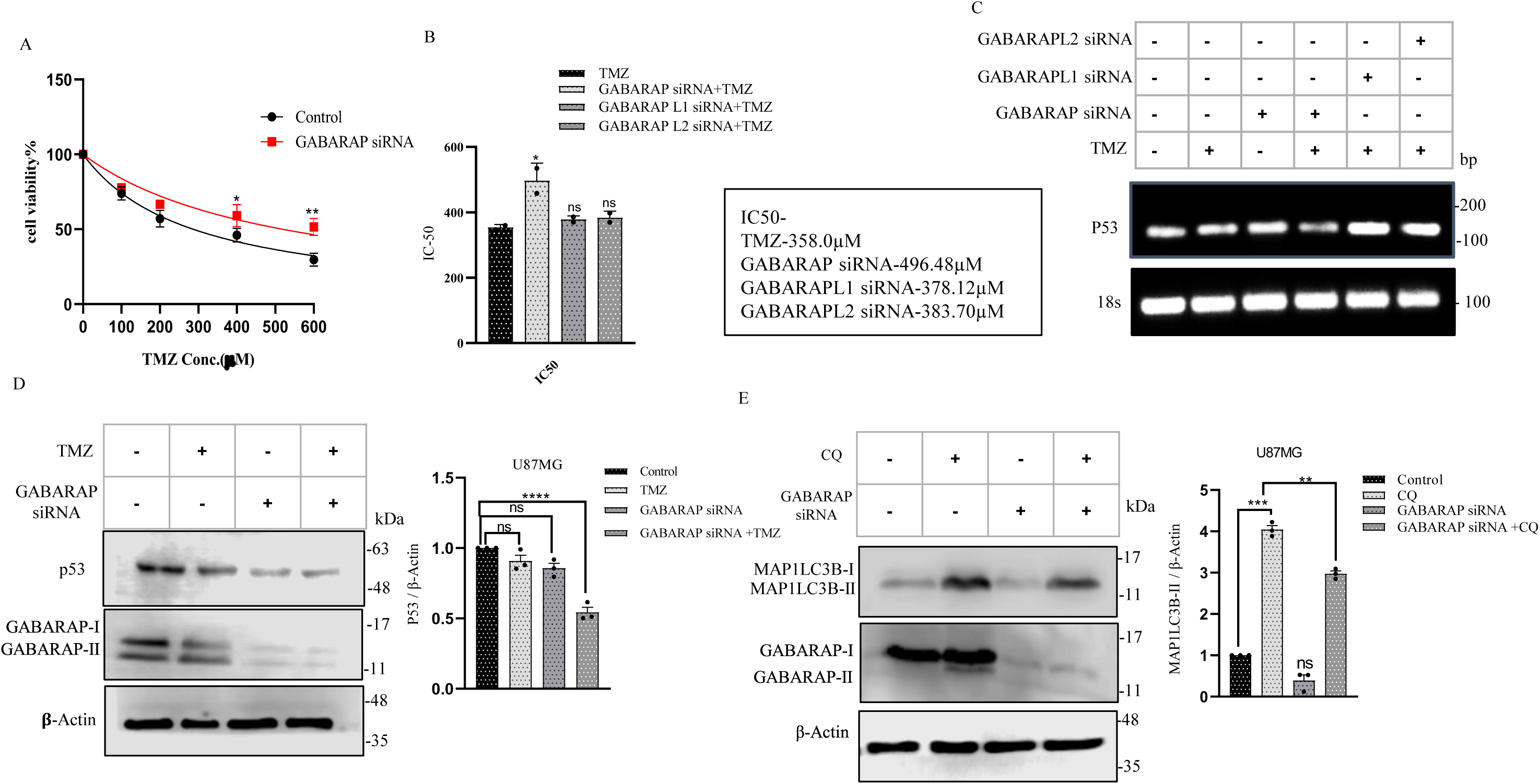
Effect of GABARAP knockdown on cell viability and expression of p53 (A) Cell viability assay for calculation of IC50 of TMZ after GABARAP knockdown (B) TMZ IC50 in Con and after knockdown of GABARAP, GABARAPL1, or GABARAPL2. (C) mRNA expression of p53 after TMZ treatment along with knockdown of GABARAP, GABARAPL1, or GABARAPL2 in U87MG cell line. (D) Western Blot of p53 and GABARAP in U87MG cell line treated with TMZ alone GABARAPsi and TMZ along with GABARAPsi. β-actin used as loading control. (E) Western Blot of MAP1LC3B and GABARAP in U87MG cell line treated with CQ alone, GABARAPsi and CQ along with GABARAPsi. β-actin used as loading control. All the statistical values have been calculated using one-way ANOVA (* p< 0.05; **p <0.01, ***p <0.001, ****p <0.0001).

### 6. Overexpression of GABARAP enhances TMZ sensitivity and increases p53 expression

Next, we wanted to evaluate the effect of GABARAP overexpression on TMZ sensitivity and p53 expression. To do so we expressed pEGFP-GABARAP in U87MG cells and observed a significant decrease in IC50 of TMZ from 358.0μM to 109.0μM (Fig 6A). This implies that overexpression of GABARAP stimulates the TMZ cytotoxicity, which might be required for suppressing glial cell proliferation. However, this synergistic effect was not observed with overexpression of GABARAPL1 and GABARAPL2 (Supp. Fig. 6A). Further, the treatment with IC50 TMZ and a combination of GABARAP overexpression with TMZ in U87 cells led to suppression of cell proliferation as shown by colony forming assay (Fig 6B). Also, the level of p53 was significantly increased after GABARAP overexpression and TMZ treatment at mRNA level (Fig. 6C) as well as at protein level (Fig. 6D). The role of GABARAP in stimulating p53 upon TMZ treatment might be essential for cytotoxic effects of TMZ as we observed that only GABARAP shows a positive correlation with p53 in brain cancer but GABARAPL1 shows a negative correlation and GABARAPL2 does not show any correlation with p53 (Supp Fig 6B, 6C, 6D). We also analyzed similarly correlated genes between GABARAP and P53 (Supp Fig. 6E) and observed 30 topmost genes, which show a correlation with GABARAP and P53. Out of these, 15 genes were positively correlated, and 15 genes were negatively correlated. Polyadenylate-binding cytoplasmic 1 (PABPC1) gene was correlated with tumor progression and diagnosis. PABPC1 is a cytoplasmic RNA binding protein component, that was broadly distributed in the cytoplasm as a regulator of mRNA stability, post-mRNA translation control, and mRNA decay. Higher expression of PABPC1 indirectly inhibited the progression, migration, and invasion of GBM cells. GABARAP was crucial in mediating TMZ cytotoxic effects via modulating p53 levels (Supp Fig 6F). Moreover, knockdown of p53 abolished the effect of GABARAP overexpression in enhancing TMZ sensitivity (Supp Fig 6G), but MAP1LC3A overexpression did not change TMZ sensitivity significantly (Supp Fig 6H).

**Figure 6.**
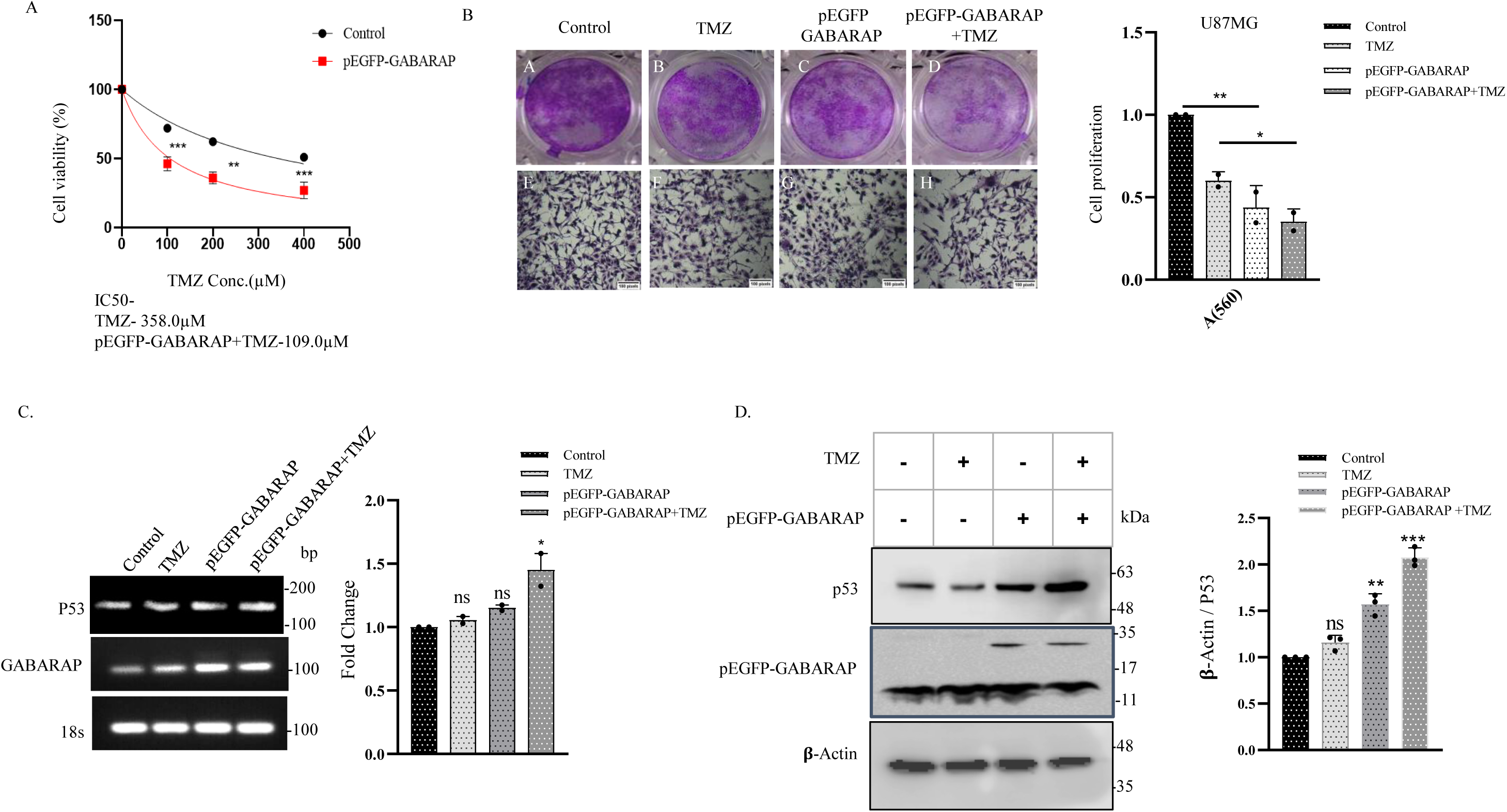
Overexpression of GABARAP enhances temozolomide sensitivity by increasing p53 expression in U87MG cell line (A) MTT assay to assess cell viability after GABARAP overexpression and TMZ treatment at different concentrations (B) Colony formation assay to assess the effect of GABARAP overexpression and TMZ treatment on cell proliferation. (C) mRNA expression analysis of p53 and GABARAP after TMZ treatment and with GABARAP overexpression (D) Western blot analysis to detect the expression of p53 and GABARAP after TMZ treatment and with GABARAP overexpression. β-actin was used as loading control. All the statistical values have been calculated using one-way ANOVA (* p< 0.05; **p <0.01, ***p <0.001, ****p <0.0001). For the IC50 calculations, non-linear regression curves were used.

### 7. Combined treatment of TMZ with CQ increases GABARAP and p53 expression and enhances glial cell death

We further wanted to analyze the effect of the combined treatment of TMZ and CQ on the expression pattern and cell viability. The dual role of autophagy in cancer is reported in GBM. Previous studies have shown that during early stages of cancer progression, the autophagy induction by mTOR inhibition contributes to suppression of cell proliferation and increased sensitivity to drug treatment in GBM for prolonged survival time^55^. Recent studies have revealed that the combination of autophagy inhibitors with chemotherapeutic drugs enhances the cytotoxicity of chemotherapeutic drugs^56^. We analyzed the effect of TMZ and CQ alone v/s combined treatment of TMZ (IC50) and CQ (IC50) for 24hours on U87MG cell viability.After combined treatment GABARAP and p53 gene expression was checked in U87MG and LN229 cells (Supp. Fig 7A, 7B). We observed that combined treatment promoted glial cell death compared to alone treatment with TMZ and CQ (Fig. 7A, 7B, Supp. Fig. 7C, 7D). The mRNA expression of both p53 and GABARAP was increased significantly in the combined treatment group (Supp. Fig. 7A, 7B). At the protein level, we observed increased expression of GABARAP-I in CQ treatment where as increased accumulation of GABARAP-II in TMZ treatment and GABARAP-I and II were increased in combined treatment (Fig. 7C, 7D, 7E). P53 protein was significantly increased in the combined treatment group, thus signifying the effect of increased GABARAP expression on enhancing or stabilizing p53 protein levels. We already found that GABARAP overexpression promoted the sensitivity of TMZ by enhancing p53 expression since the knockdown of p53 abrogated the GABARAP induced reduction in TMZ IC50. Our study thus revealed the crucial GABARAP-p53 axis that is involved in promoting the cytotoxic effect of TMZ and hence it is probable that GBM patients with high expression of GABARAP might have a better therapeutic response to TMZ treatment.

**Figure 7.**
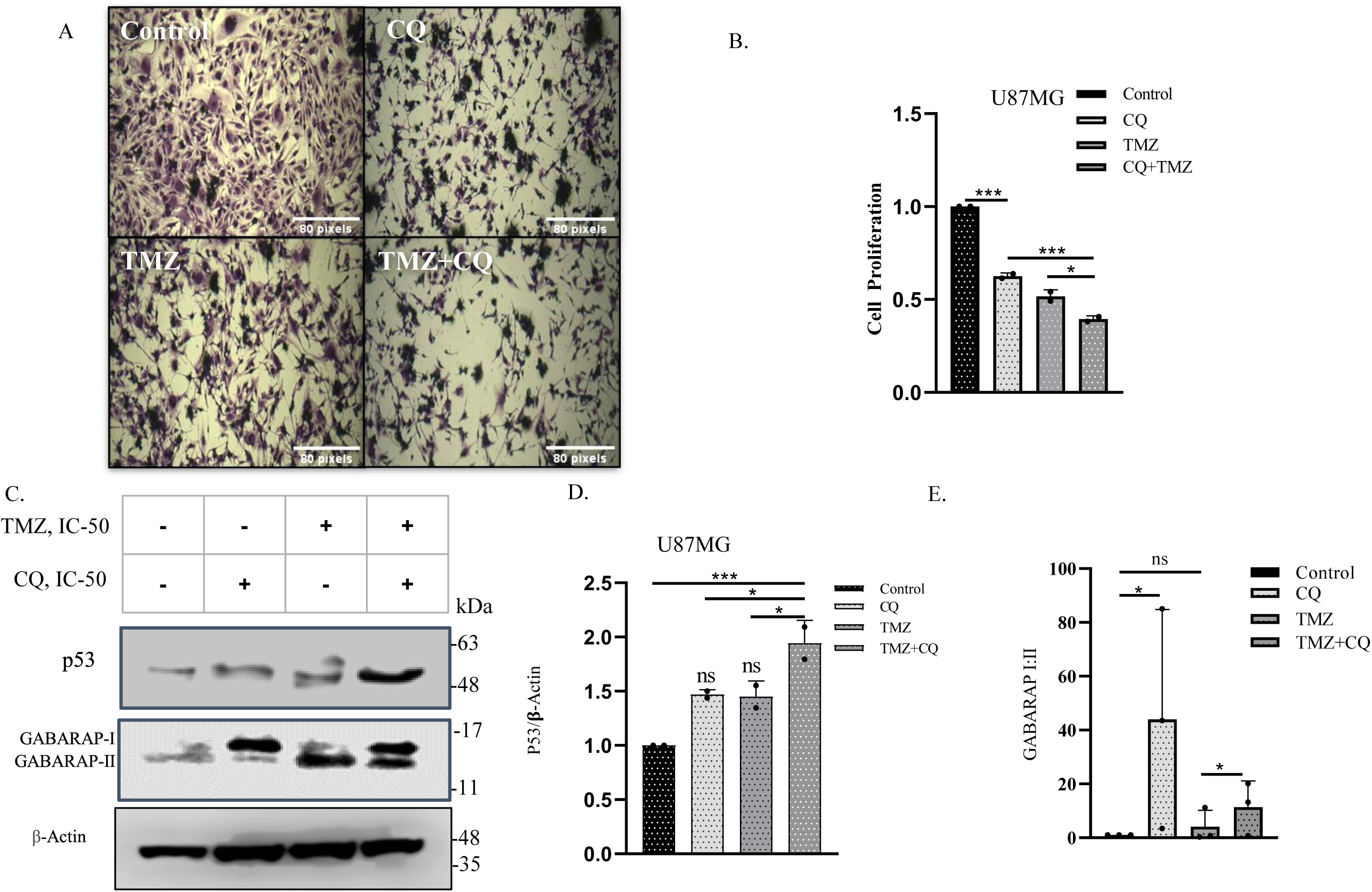
Combination treatment of TMZ and CQ (A-B) Cell images and MTT assay after the 24hours treatment of TMZ(IC50), CQ(IC50), and a combined treatment of TMZ and CQ in U87MG. (C-D) Western blot analysis was used to detect the expression of GABARAP and p53 in the U87MG cell line after TMZ(IC50), CQ(IC50), and combined treatment for 24 hours, β-actin was used as loading control. (E) Non-lipidated (I) and lipidated (II) forms of GABARAP were quantified after CQ, TMZ and combined treatment of CQ with TMZ. All the statistical values have been calculated using one-way ANOVA (* p< 0.05; **p <0.01, ***p <0.001, ****p <0.0001).

## Discussion

Glioblastoma is the most aggressive and prominent cancer because of its hypermutant behavior. Somehow, the reason for the low survival rate is due to the resistance of many anticancer therapies like chemotherapy and radiotherapy^41–43^. Several studies have demonstrated the cytoprotective and pro-survival functions of autophagy, allowing cancer cells to survive the stressful conditions in the tumor microenvironment^60–63^. Inhibition and modulation of autophagy and apoptosis pathways is a major contributing factor in TMZ chemoresistance^64,65^. Therefore it is necessary to understand the precise mechanism of autophagy modulation to improve the therapeutic efficiency of TMZ. Autophagy inhibition is a promising strategy since inhibiting autophagy promotes cell death and improves TMZ sensitivity in GBM cells ^53,66,67^. However, the role of ATG8 specially GABARAPs has not been elucidated with respect to TMZ sensitivity ^68^. CQ has been previously explained in a study that it transiently affects lysosome pH and thus only blocks autophagosome-lysosome fusion, not the degradative property of lysosomes^69^. However, it also has been seen that CQ simultaneously disorganizes Golgi/endosomal complexes which might occur as side effects causing problems in results interpretation. Identifying inhibitors other than antimalarial drug CQ that are used in combination with standard therapy to increase the survival probability of GBM patients, might have better therapeutic relevance. A previous study demonstrated that the mutagenic property of TMZ causes hypermutations in GBM that lead to form a clone and provide resistance against therapy^40,65^.

Our finding demonstrates that overexpression of GABARAP enhances the cytotoxicity of TMZ. In this study, we found that GABARAP is highly expressed in most GBM and LGG patients as compared to other ATG8 genes^6^. Moreover, GABARAP was positively correlated, while MAP1LC3A was negatively correlated with overall patient survival. Further in-vitro analysis in glial cell lines demonstrated that autophagy inhibition and induction differentially impacted GABARAP family member expression signifying member-specific functions. To ascertain the role of individual members, their non-lipidated levels were quantified, which corroborated well with the changes in the levels of their transcripts. This might imply that cells prefer to use the pool of GABARAP family members in a sequential way where GABARAPL1 and GABARAPL2 are preferably lipidated upon CQ treatment, and GABARAP total protein was increased to take care of lipidation at later stages. Unique functions of GABARAP are previously reported^70,71^. Various findings analyzed that autophagy is a major contributor to TMZ resistance^72^, but still, the involvement of autophagy-related genes and its molecular mechanism during therapy remains unclear. Considering the differential response of GABARAPs we reasoned whether TMZ might affect their expression and observed enhanced expression of GABARAP in a dose and time dependent manner, while MAP1LC3A expression was decreased significantly. GABARAP has previously been associated with inducing cytotoxicity of other chemotherapeutic drugs^73–75^. Accordingly, the knockdown of GABARAP significantly increased the TMZ IC50 while the knockdown of GABARAPL1 and GABARAPL2 did not (Fig 5B). Moreover, TMZ treatment along with GABARAP knockdown led to a significant decrease in p53 levels at mRNA as well as protein level, which was not observed, with knockdown of GABARAPL1 and GABARAPL2.

The essential roles of GABARAP family members in autophagy have been observed previously ^68,76^. To validate if GABARAP knockdown also perturbs the autophagy process we checked LC3B levels upon GABARAP knockdown and CQ treatment and indeed there was reduced level of LC3B upon GABARAP knockdown (Fig 5E). Interestingly, GABARAP overexpression led to a decrease in TMZ IC50 and caused an increased expression of p53 upon TMZ treatment, thus increasing TMZ sensitivity by affecting p53 expression. Finally, we performed a combined treatment of CQ and TMZ and observed a synergistic effect in promoting cell death. Interestingly, while CQ treatment predominantly increased GABARAP-I levels, TMZ treatment enhanced GABARAP-II levels, and combined treatment led to overall drastic increase in GABARAP levels. Moreover, combined treatment with TMZ and CQ enhanced GABARAP significantly and resulted in synergistic increase in p53 expression^77^. These results indicate that GABARAP is related to the survival and proliferation of the GBM cells and GABARAP can be used as a good prognostic marker of GBM. The GBM samples with high expression of GABARAP would possibly respond better to TMZ as well as combination therapy of CQ and TMZ. In this study, we explored the differential expression of GABARAP family members in response to CQ and TMZ and observed that family members behave differently in response to different stresses. Therefore, the cancer cells utilize the individual GABARAP family members to adapt better for chemotherapy and other stress conditions and, this subtle regulation can be explored further to acquire better therapeutic advantage in GBM and also in other cancers.

## Methods

### 1. Patient Samples

Surgically resected GBM tumor tissues were collected from the Department of Neurosurgery, AIIMS, Delhi, after due consent from patients and ethical clearance from the institute ethics committee (<u>IEC-711/07.08.2020, RP-43/2020</u>). Tumor tissues were collected in RNA-Later solution (Ambion, USA, catalog no. AM7020), incubated at 4°C for 24 hours, and shifted to −70°C after removal of RNA-Later solution, until further use. The histopathological diagnosis of tumors was done by neuropathologist Prof. Chitra Sarkar/Prof. Vaishali Suri. Total RNA from frozen tumor samples was extracted using TRI Reagent (Sigma-Aldrich, USA) and quantified using NanoDrop ND-1000 spectrophotometer (Thermo Fisher Scientific, USA). Genomic DNA was removed by DNase I (MBI Fermentas, Hanover, MD) treatment. Reverse transcriptase reactions were performed using random decamers (MWG, India) and reverse transcriptase enzyme (MBI Fermentas, Hanover, MD) in 20μl reaction volume. Normal human brain total-RNA (Clontech, USA, catalog number 636530) was used as the control for q-PCR analysis.

### 2. In- silico analysis

For our analysis, we initially selected user-friendly and interactive web server that helps us to analyze our gene set. In total three datasets were used in present study. GEPIA database was used to check the expression of all the genes in two different stages GBM and LGG. For all the datasets, a comparison between TCGA normal and GTEx data were used. Parameters such as Log_2_FC value and p-value cutoff was set to 1.00 and 0.01, respectively. Further PCA plot analysis was performed to check the connection between these genes in both conditions. Survival analysis of GBM was calculated by using Glioma Bio Discovery Portal (Glioma-BioDP) database^78^. The output from BioDP database, the aggregates come in two sets: a complete one with expression data for every gene that is featured in all platforms (Aggregates) and filter set (Verhaak core) which contains 840 genes that were stipulated by Verhaak et.al., 2010 to have special significance for characterizing the molecular subtype of GBM.

### 3. Cell culture and reagents

Glioblastoma multiform cell lines (U87MG, LN229) were purchased from the National Center for Cell Science (NCCS), Pune, India. The cells were cultured in 90% Dulbecco’s modified Eagle’s medium (DMEM) (AL007F-500ML, Himedia) with 10% Fetal bovine serum (FBS) (RM10409, Himedia, Heat inactivated) and 1% penicillin-streptomycin (A001-100ML Himedia) at 37°C in a humidified atmosphere with 5% CO_2_. During drug treatment, the cell culture medium was switched to DMEM without FBS in the case of chloroquine treatment and with serum media in Temozolomide (T2744, TCI) treatment. The chloroquine drug (10mM) dissolved in Phosphate buffer saline (PBS) (TS1101-10X1L, Himedia), and TMZ (100mM) dissolved in dimethyl sulfoxide (DMSO) (TC185, Himedia) and stored at −20°С.

### 4. RNA isolation and Real-time PCR

To measure gene expression, U87MG, and LN229 cells were seeded in six-well plates at 3×10^5^ cells/well. Total RNA was isolated by using the TRI Reagent (Sigma-Aldrich, USA, T9424) according to the manufacturer’s protocol. The concentration of RNA and its quality was measured using NanoDrop. For cDNA synthesis, 1000 ng of RNA was used per sample. Verso cDNA synthesis kit (Thermo Fisher Scientific, K1632) was used according to the manufacturer’s guidelines using random hexamer. A total of 20µl reaction was set up for cDNA synthesis. The reaction was finally diluted to 1:5 in nuclease-free water and 2.5µl of it was used per reaction of qPCR. For quantification of transcript levels, SYBR green-based qPCR was performed using the SYBR™ Green PCR Master Mix (Thermo Fisher Scientific, 4309155). The qPCR was initiated with 2 min at 50°C, followed by denaturation of 10 min at 95°C. This was followed by 35 cycles of 15 s at 95°C, 60 s at annealing temperature (57-60°C) and 20 s at 72°C. Relative expression of the analyzed genes were calculated compared to the housekeeping gene *18s* by using the 2^−ΔΔ*C*^T-method. Three independent experiments in triplicate were performed for each condition. Gene-specific primers were designed and are listed in Table 3. Supp. Fig 9 shows the exact amplicon size of primers with the help of 100bp ladder.

### 5. Cell viability assay

Cell viability was measured by 3-(4,5 dimethylthiazol-2-yl)-2.5-diphenyl-^2^H-tetrazolium bromide (MTT) (MB186-1G, Himedia) assay. MTT assay is the fastest and most precise method to analyze cell growth. In brief, cells were seeded in 96-well microplates at 3.5 x 10^3^cells/well. The tumor cells were treated using different concentrations of the drugs for 48 to 72 hours. We set up a control group and a zero-adjustment group. At specified time points, 10ul of MTT solution (5 mg/mL) was added and incubated an additional 4 hours at 37°C before the end of the incubation duration; the reaction was stopped by the addition of 100 μL dimethylsulfoxide (DMSO) (TC185, Himedia). The optical density (OD) or absorbance was read at the 540-570 nm range.

### 6. Colony formation assay

To study the effect of different drugs like temozolomide and chloroquine after GABARAP overexpression and siRNA transfection, a colony formation assay was performed in the U87MG cells. Cells (1000/per well) were seeded onto 24 well culture plates in DMEM media with 10% FBS. The cells were treated with CQ and TMZ drugs and incubated for 24-48 hours at 37°С in a 5% CO2 incubator. After colonies appeared, the cells were fixed with 4% paraformaldehyde solution for 10 min and stained with 1% crystal violet (TC510-25G, Himedia) stain. These cells were then washed twice with PBS. The captured stain was taken using 0.1% SDS solution, and absorbance was measured at 570nm.

### 7. Western Blotting

Cells were washed with phosphate-buffered saline (PBS) and then lysed in NP40 lysis buffer (25 mM Tris pH 8, 150 mM NaCl, 0.1% NP-40, 0.1 mM EDTA) containing 1X protease inhibitor cocktail (Himedia, ML051-1ML), for 30 min on ice. Total protein concentrations were analyzed by using a BCA kit (Micro BCA^TM^ Protein Assay kit 23235 Thermo Fisher). The lysates were denatured in 2X SDS Sample buffer at 95°C for 5 mins and run on 15% tris-glycine gel. We used 50ug proteins/well and loaded on SDS-PAGE, followed by transfer onto a PVDF membrane (Bio-Rad 1620177, Pore size .2µm) at 65V. To avoid the non-specific binding sites, membranes were incubated for 1 hour at room temperature with 5% skim milk in TBS and 0.1% tween-20, pH=7.5 (TBS-T) and later membranes were gently washed with TBST three times. These blots were incubated with (1:1000) dilutions of primary antibody (In 1xTBST) for overnight at 4°C on rocker shaker. Next day, after three gentle washing with TBST, appropriate secondary antibodies (Anti-mouse or Anti-rabbit) with a dilution of (1:3000) were added at room temperature for 2 hours. The blots were then processed for visualization using an ECL (Enhanced Chemiluminescence) (Thermo Scientific, Product no: 32106) substrate, and the images were taken on the blot scanner (Licor, c-digit). The antibody used was: Rabbit-monoclonal antibody LC3A (1:1000, D50G8, Cell Signaling), rabbit monoclonal antibody LC3A/B (1:1000, D3U4C, Cell Signaling), Mouse monoclonal antibody Beta-actin (1:1000,8H10D10, Cell signaling), Anti-mouse IgG HRP linked antibody (7076P2,1:5000, Cell Signaling), Rabbit monoclonal GABARAP antibody(1:1000, 13733 (E1J4E) Cell Signaling), Rabbit-monoclonal antibody GABARAPL1 (1:1000, D5R9Y(26632), Cell Signaling), Rabbit-monoclonal antibody GABARAPL2 (1:1000, D1W9T(14256), Cell Signaling). Rabbit-monoclonal antibody p62 (1:1000, 39749, cell signaling) Cell, Rabbit-monoclonal antibody LC3B (1:1000, 3868, cell signaling). We used 50μg protein and individually probed for all three GABARAP family members. We then used one of the blots and reprobed for β-actin. We performed the stripping using the mild stripping buffer (Glycine,15; SDS, 1g; tween20, 10ml; Deionized water, 800 ml; pH to 2.2 with HCL), followed by washing the membrane 3 times with 1x TBST. Further the blot was blocked using 3% BSA and reprobed for β-actin. The densitometry analysis was done using Image J software.

### 8. siRNA Transfection and Plasmid transfection

To knock down the GABARAP, GABARAP L1, and GABARAP L2, U87MG cells were transfected with three different siRNAs against GABARAP (NM_0072782-AS), GABARAPL1 (NM_0314124-AS) and GABARAPL2 (NM_0072857) at 60% - 80% confluency by using Lipofectamine 3000 (Thermo Fisher Scientific, L3000008) according to the mentioned protocol. For the non-targeting siRNA, we have used Scrambled control siRNA from GeneX India (catalog no SR-CL000-005). The following amount of Lipofectamine 3000 was used per well: 4µl/6-well, 2µl/24-well. Cells were transfected with 20 to 50 nM of siRNA in only DMEM. After 24 hours, the media was changed and cells were re-transfected with the same amount of siRNA and left for 24 hours. After a total of 48 hours, the RNA was isolated and Real-time PCR was performed. For validation, the PCR sample was run on the agarose gel. Cells were transfected with pEGFP-hGABARAP Plasmid (Addgene, 87871) for overexpression by using Lipofectamine 3000 according to manufacturer’s protocol. Plasmids pEYFP-hLC3A, pEYFP-GABARAP, pEYFP-GABARAPL1 and pEYFP-GABARAPL2 were kindly obtained from Dr. Silke Hoffmann, Julich, Heinrich Heine University, Germany. Briefly, 3,00,000 cells were seeded in a 6-well plate and cells were transfected with 1ug plasmid DNA and 3ul Lipofectamine 3000 in fresh medium and pEGFP (addgene 148304) was used as a control. Plasmid transfection was done for 24 hours and after 24 hours of incubation cells were processed for RNA and protein isolation. To monitor the autophagy flux, we used double tagged LC3B protein. We transfected the LN229 cells with mCherry-GFP-LC3B plasmid (Addgene, 170466) followed by treatment with TMZ for 24 hours and Confocal imaging of mCherry-GFP-LC3B vesicles.

### 9. Statistical Analysis

All statistical analysis was done using GraphPad Prism 8.4.0. Student’s t-test was used for analyzing the expression in patient samples. One-way ANOVA with multiple comparisons test was used to estimate the difference between groups. A p-value less than 0.05 is considered statistically significant (* = p<0.05, ** = p<0.01, *** = p<0.001, **** = p<0.0001). All experiments were performed in triplicate. All the densitometric calculations were done using ImageJ.

## Abbreviations

GBM: Glioblastoma Multiforme
LGG: Low Grade Glioma
CQ: Chloroquine
HCQ: Hydroxychloroquine
BafA1: Bafilomycin A1
TMZ: Temozolomide
GABARAP: Gamma aminobutyric acid receptor -associated protein
GABARAPL1: Gamma aminobutyric acid receptor -associated protein like-1
GABARAPL2/GATE-16: Gamma aminobutyric acid receptor -associated protein like-2
MAP1LC3A: Microtubule associated protein 1A/1B -light chain 3 alpha
MAP1LC3B: Microtubule associated protein 1A/1B -light chain 3 beta
MAP1LC3C: Microtubule associated protein 1A/1B -light chain 3 gamma
TP53: Tumor protein P53

## Acknowledgment

We thank Dr. Silke Hoffmann, Julich, Heinrich Heine University, Germany for kindly providing the plasmids pEYFP-GABARAP, pEYFP-GABARAPL1 and pEYFP-GABARAPL2. We thank Department of Biochemistry, Central University of Rajasthan for providing a research facility and DST-SERB start-up research grant (SRG/2020/000392), ICMR extramural research Grant (52/27/2020-BIO/BMS) and SERB-Power Grant (SPG/2021/002833) for providing research funds to Dr. Bhawana Bissa.

## Credit Statement

**Megha Chaudhary:** Conceptualization, writing, Figure preparation, Experimental work; **Naveen Soni**: Figure preparation, writing, editing, Experimental work; **Shweta Dongre**: Revision experiments and proof reading; **Ashwani Tiwari**: Revision experiments and proof reading; **Nargis Malik**: Sample collection, editing; **Kunzang Chosdol:** Ethics approval, Sample collection, proof reading, editing; **Bhawana Bissa**: Conceptualization, writing, supervision

## Conflict of Interest

Authors declare no conflict of interest.

## Data Availability Statement

The data is available on Mendeley DOI: 10.17632/9zczv644bv.1

## Figure Legends

**Supp Fig. 1.**
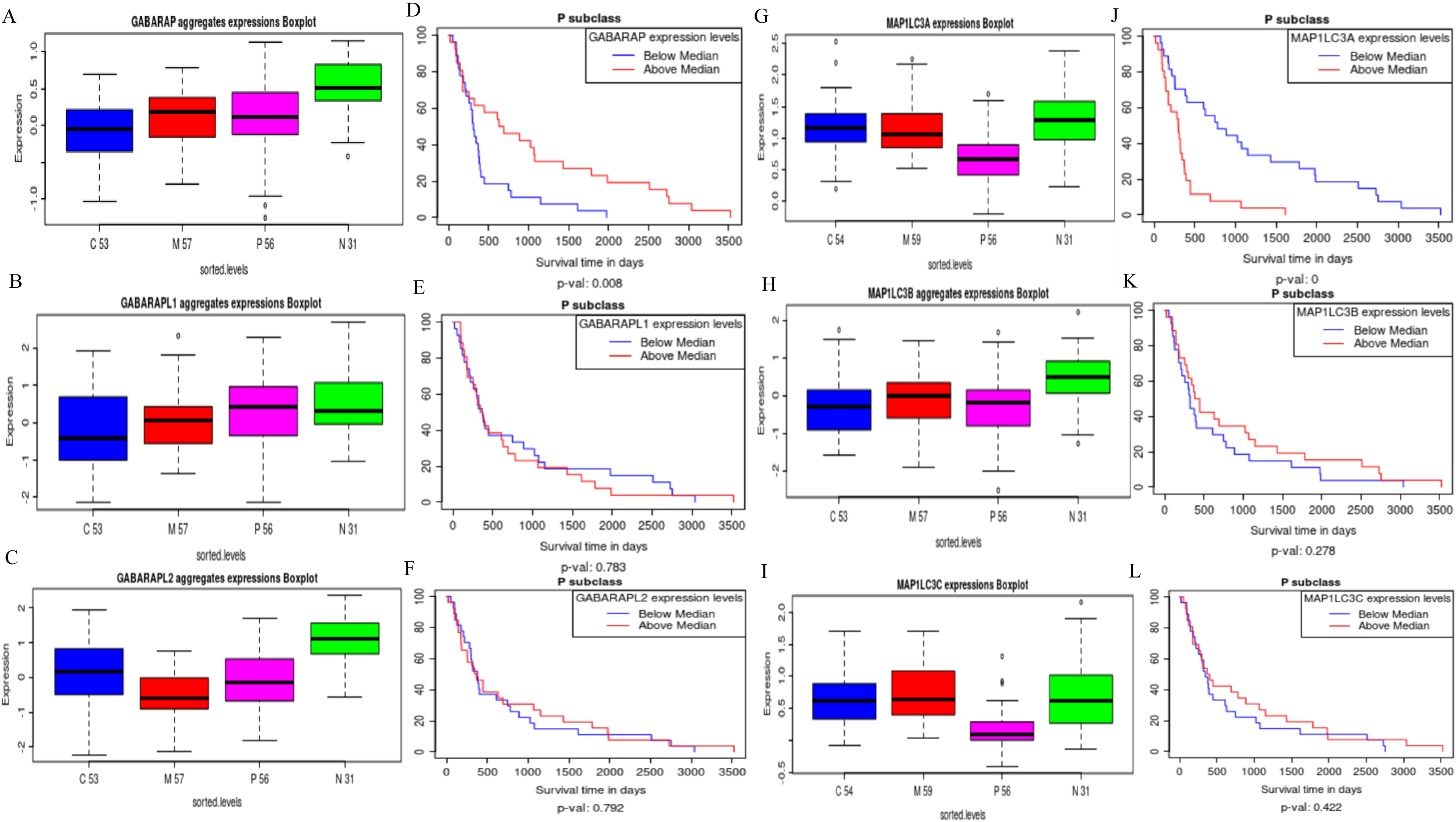
(A-C and G-I) The boxplot shows the distribution of GABARAP’s and MAP1LC3’s member gene expression according to the molecular subtype of each sample: x-axis represents the classical (C), mesenchymal (M), proneural (P), neural (N) subtypes. GABARAP, GABARAPL2, and MAP1LC3B genes were overexpressed in the neural subtype and the y-axis shows the z-score for each sample over the entire gene platform panel. (D-F and J-L) Kaplan-Meier analysis of the around 10-year survival rates of GBM patients with low and high level of all genes. (D) High expression of GABARAP promotes survival in GBM patients. (J) Low expression of MAP1LC3A enhances survival in GBM patients.

**Supp Fig. 2.**
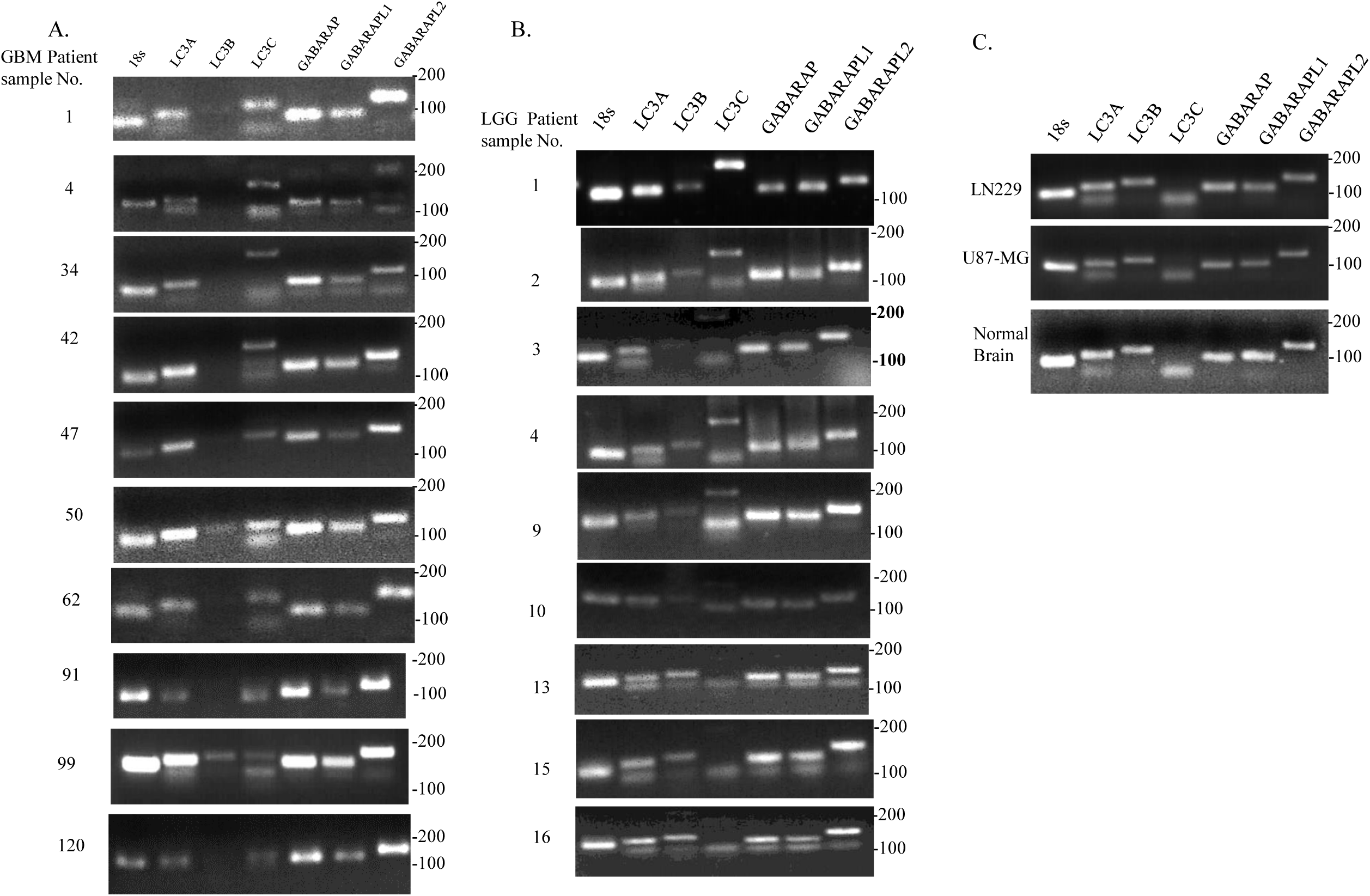
Confirmation of amplicon size of all GABARAP and MAP1LC3 family members by agarose-gel electrophoresis. (A) Agarose-gel electrophoresis showing PCR products of the expected size of genes from the quantitative RT-PCR reaction. GABARAP gene shows the highest expression in all GBM patient samples (B) Another splice variant of MAP1LC3C was amplified in maximum LGG patient samples. (C) Expression patterns were also analyzed in GBM cell lines and normal brain samples.

**Supp Fig. 3.**
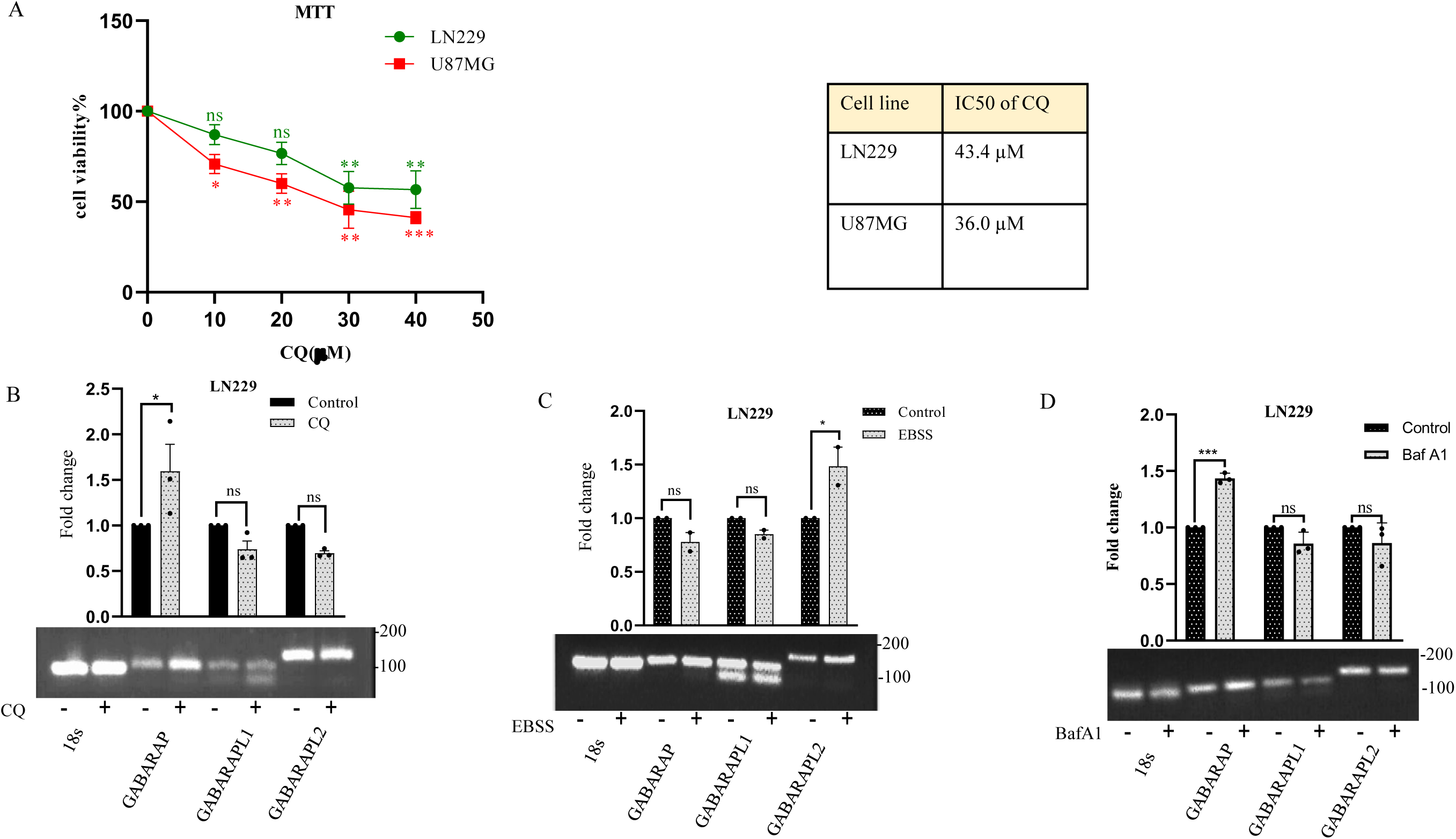
(A) Analysis of IC50 value for CQ in U87MG and LN229. (B-D) Expression analysis of GABARAP family members after CQ treatment (B) EBSS treatment (C) and BafA1 (D) treatment in the LN229 cell line. For the IC50 calculations, non-linear regression curves were used.

**Supp Fig. 4.**
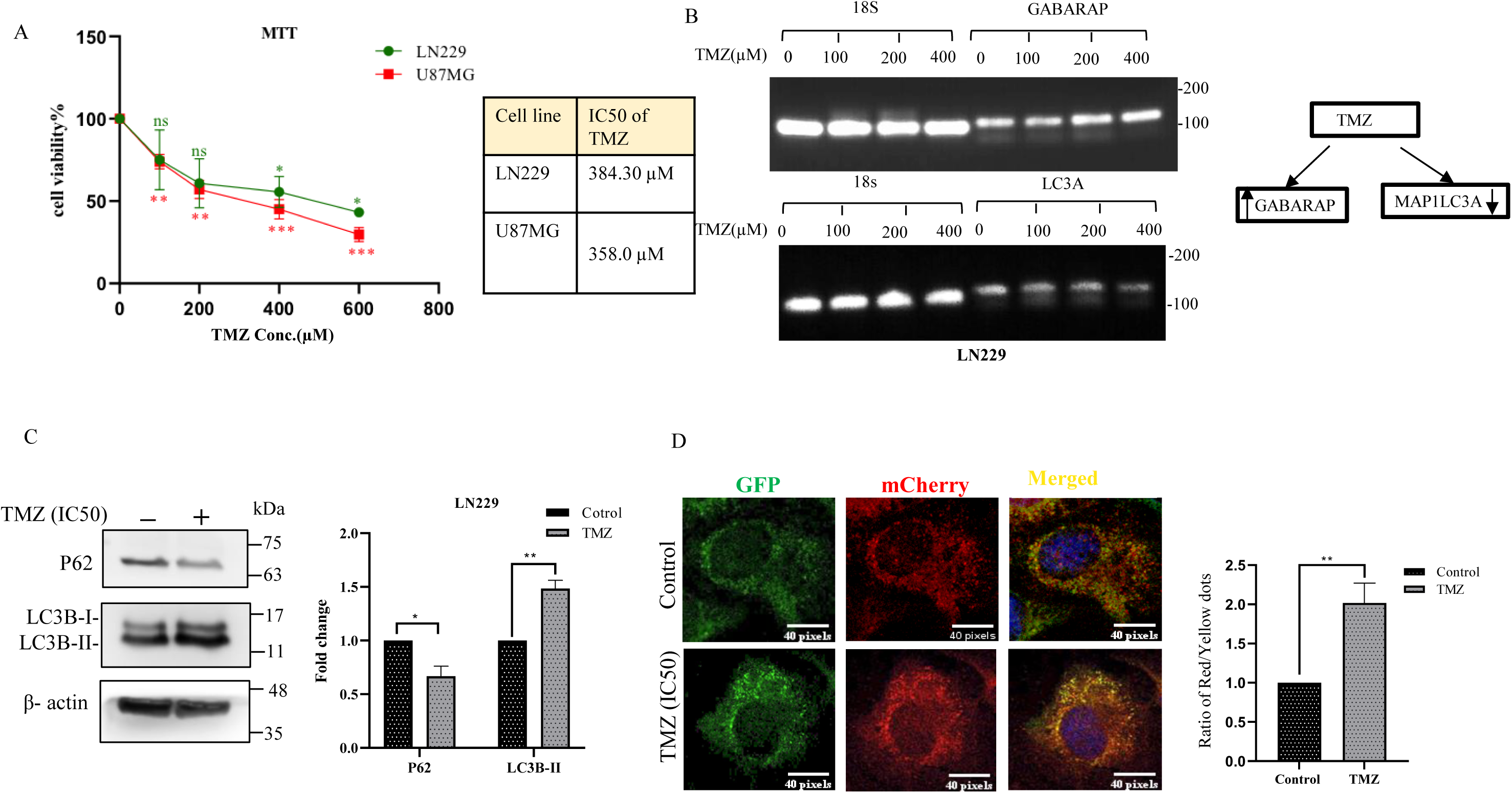
(A) Analysis of IC50 value for TMZ (100µM, 200µM, 400µM, and 600µM) in U87MG and LN229 cell line. (B) Expression analysis of GABARAP and MAP1LC3A at different concentrations of TMZ (100µM, 200µM, and 400µM). For the IC50 calculations, non-linear regression curves were used. (C) Western blot of p62 and LC3B to analyze the effect of TMZ on autophagy. β-actin was used as a loading control. (D) Confocal microscopy to demonstrate mcherry-GFP-LC3 puncta in response to TMZ treatment.

**Supp Fig. 5.**
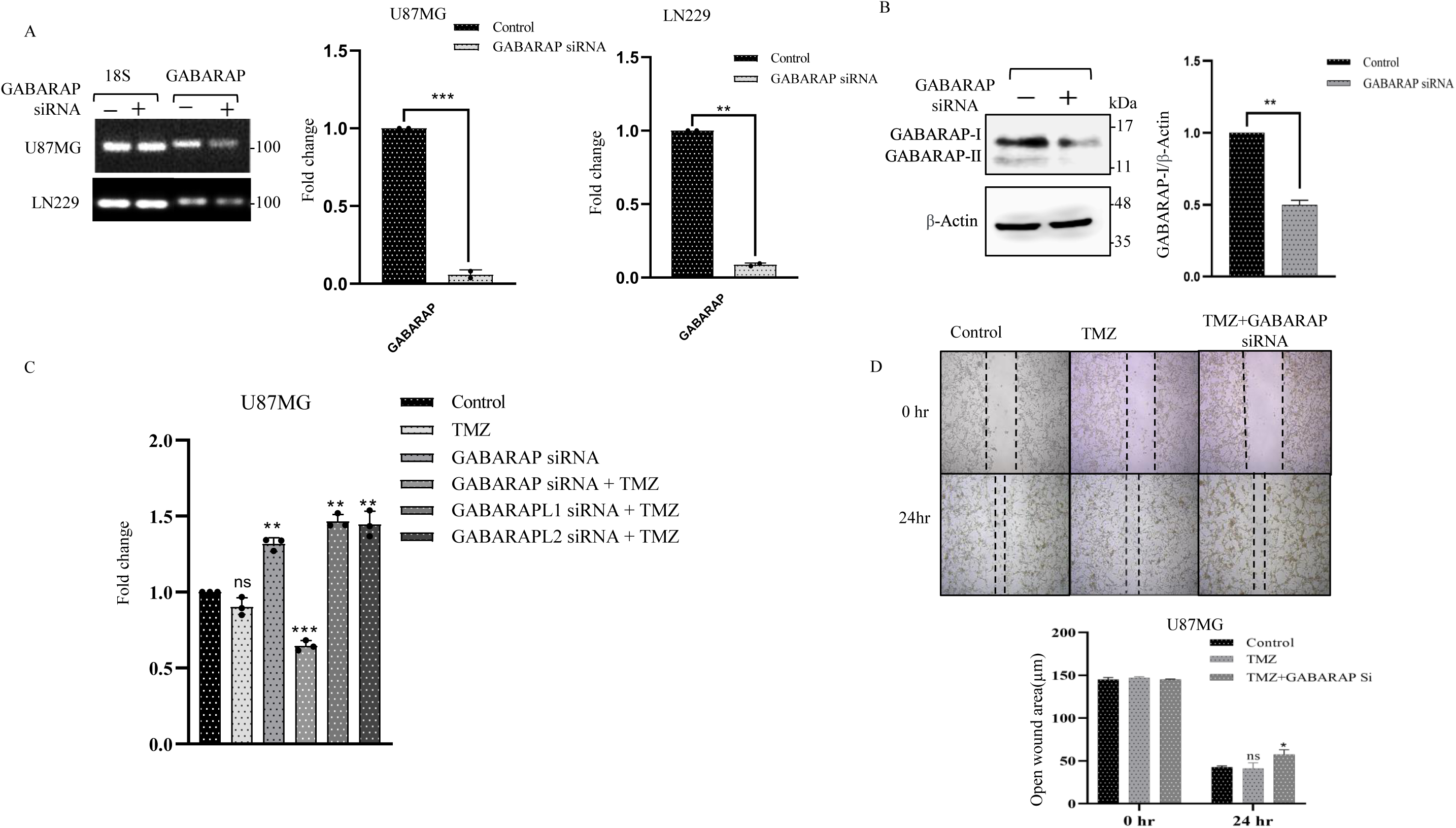
(A) A 1% agarose gel showing GABARAP knockdown in U87MG and LN229 performed by RT-PCR. 18s was used as internal control. (B) Western blot depicting GABARAP knockdown in U87MG. β-actin was used as the loading control. (C) p53 expression after TMZ treatment alone and along with knockdown of GABARAP, GABARAPL1 and GABARAPL2. (D) Migration assay in U87MG cell line for 48 hours to check the migration of cells after IC50 TMZ treatment and TMZ treatment with GABARAP knockdown.

**Supp Fig. 6.**
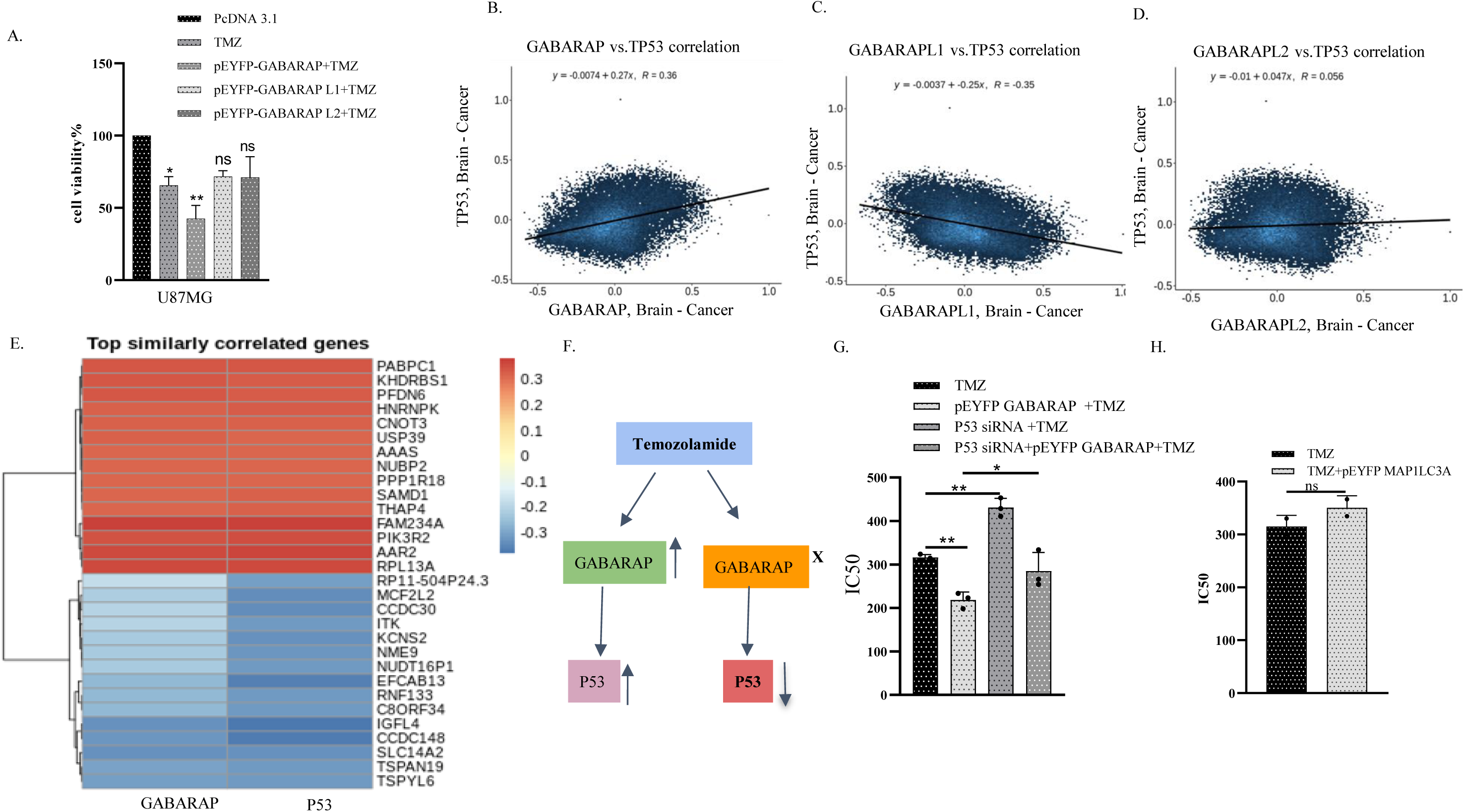
(A) MTT assay after TMZ treatment and with overexpression of pEYFP-GABARAP, pEYFP-GABARAPL1, and pEYFP-GABARAPL2. (B-D) Correlation of p53 with GABARAP, GABARAPL1, and GABARAPL2 in brain cancer. (E) Similarly correlated genes between p53 and GABARAP. (F) Schematic of the effect of TMZ on GABARAP and p53 expression. (G) IC50 graph of TMZ after GABARAP overexpression with p53 knockdown. (H) IC50 graph of TMZ after MAP1LC3A overexpression.

**Supp Fig. 7.**
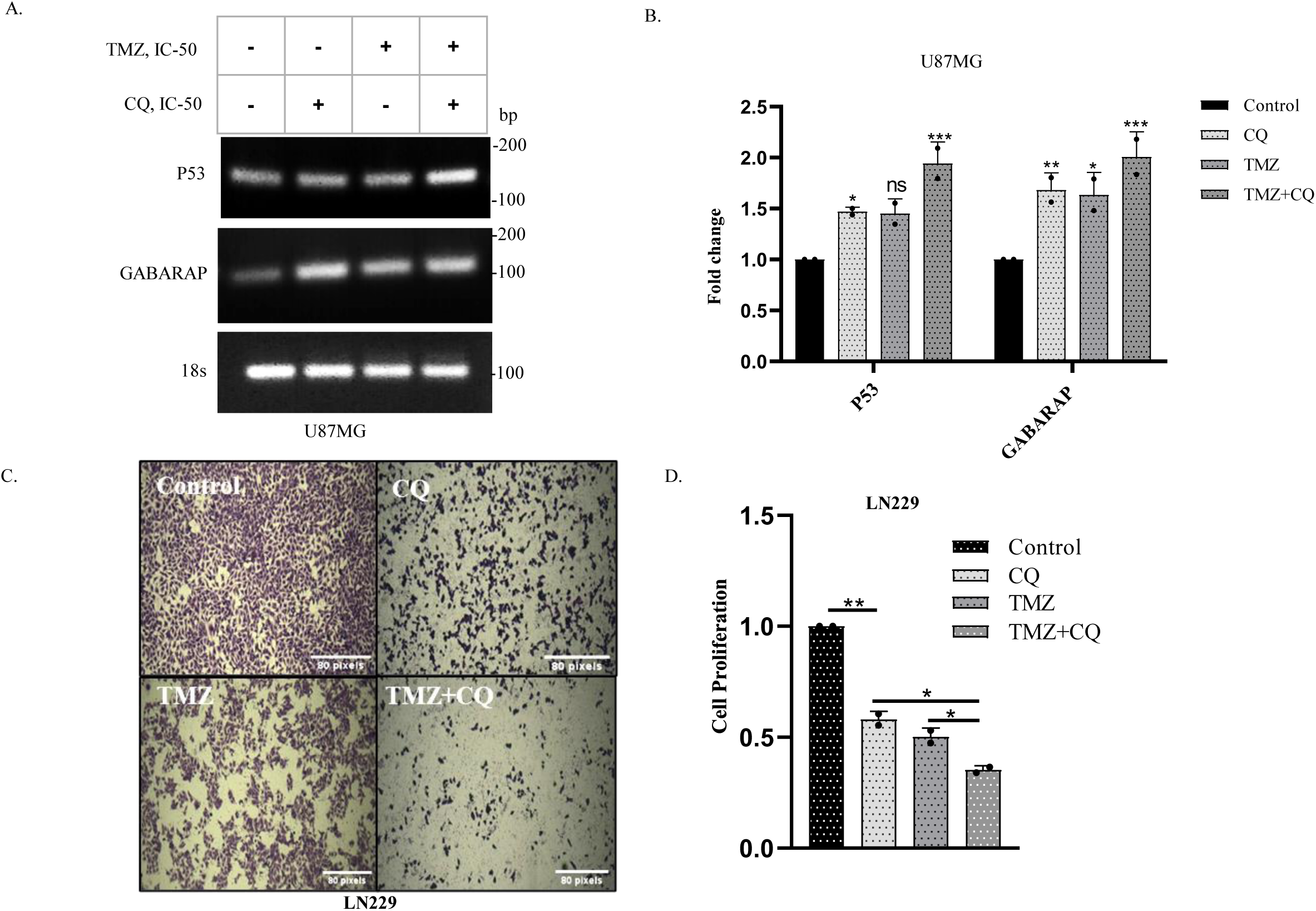
(A-B) mRNA expression analysis of p53 and GABARAP after treatment with TMZ (IC50), CQ (IC50), and combined treatment for 24 hours in U87MG cell line. 18s was used as internal control. (C-D) Cell images and cell proliferation of LN229 cell line after treatment with TMZ (IC50), CQ (IC50), and combined treatment for 24 hours for quantification of cells with crystal violet staining.

---

**Figure 3.**
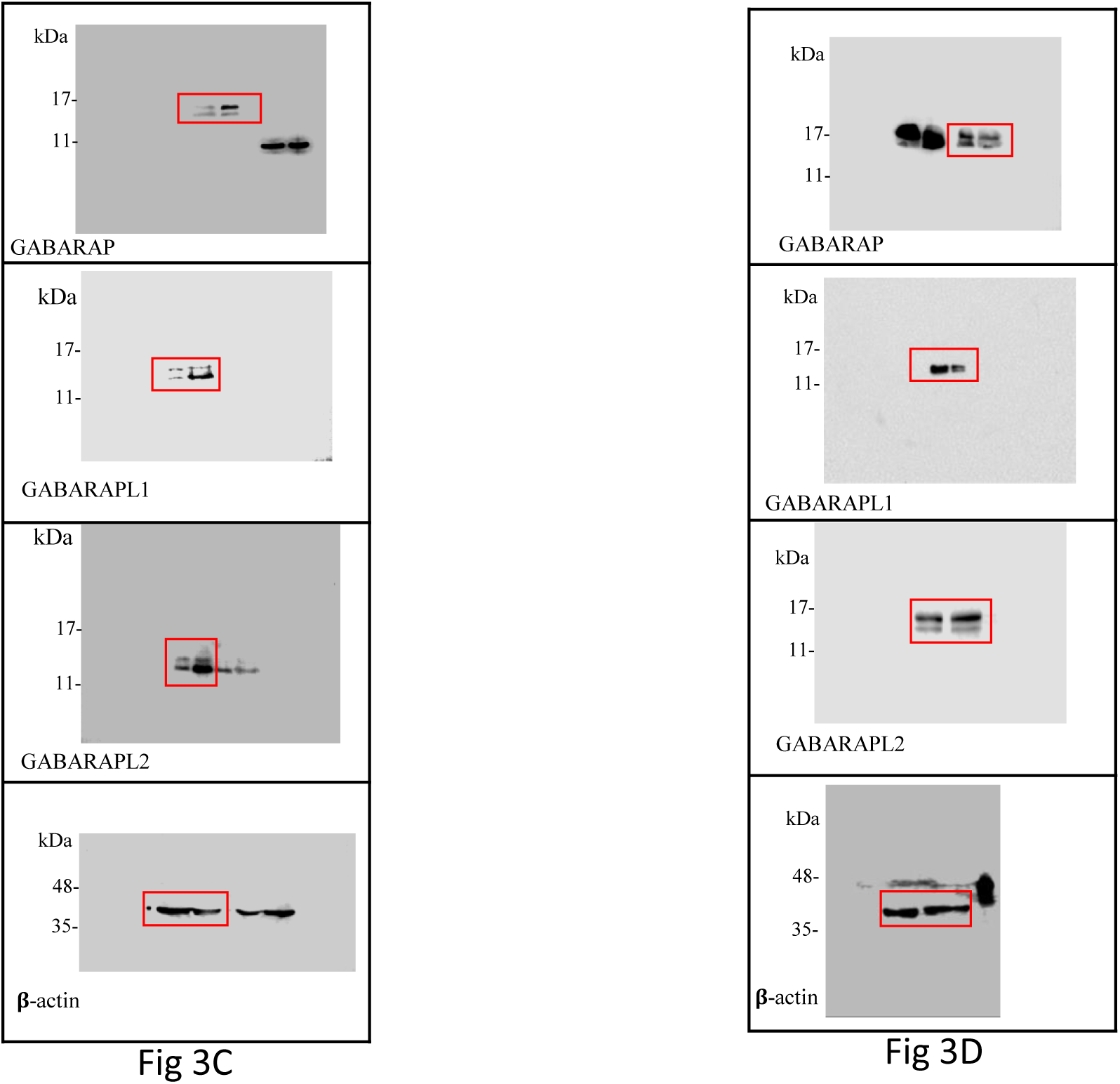

**Figure 4.**
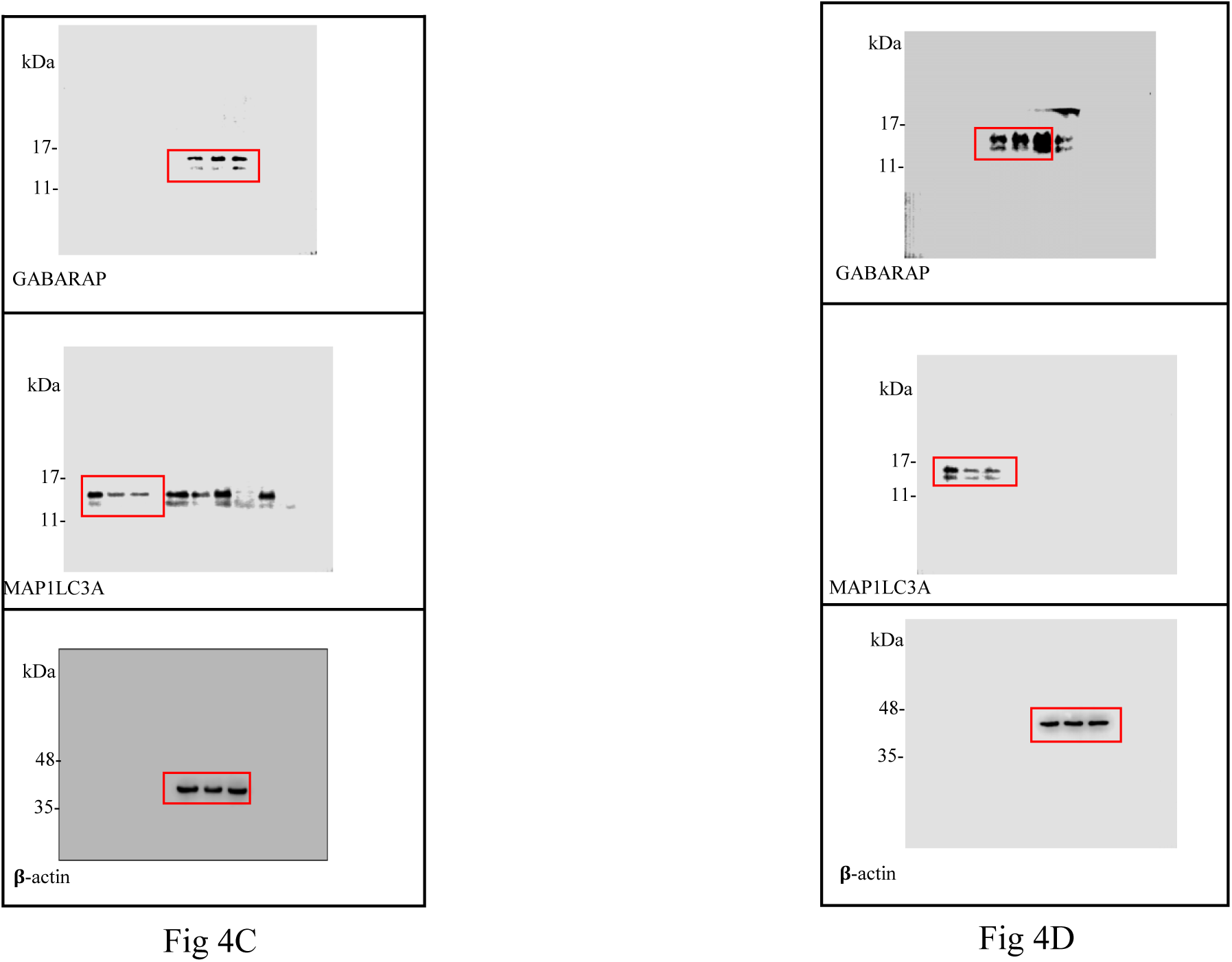

**Figure 5.**
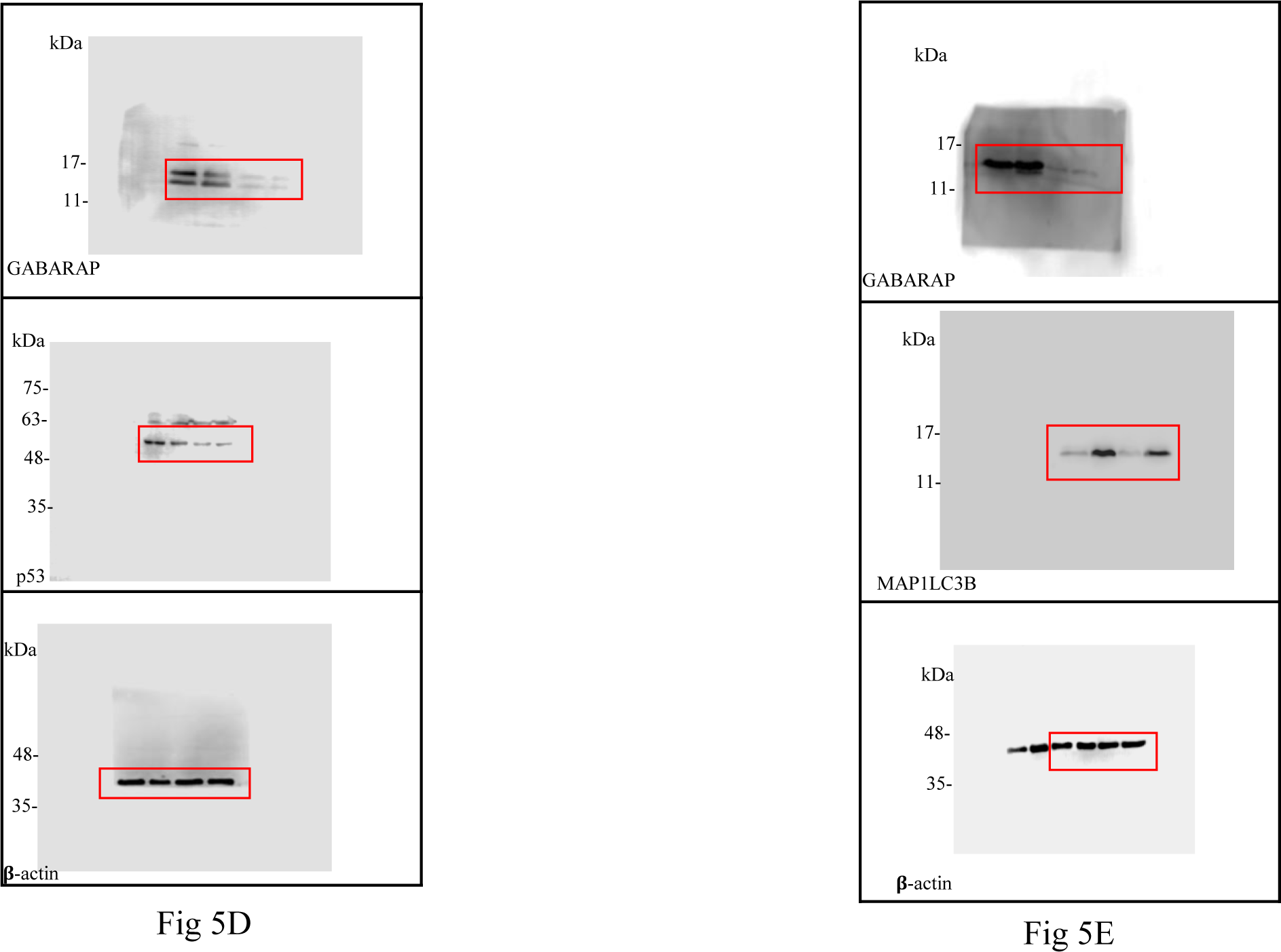

**Figure 6.**
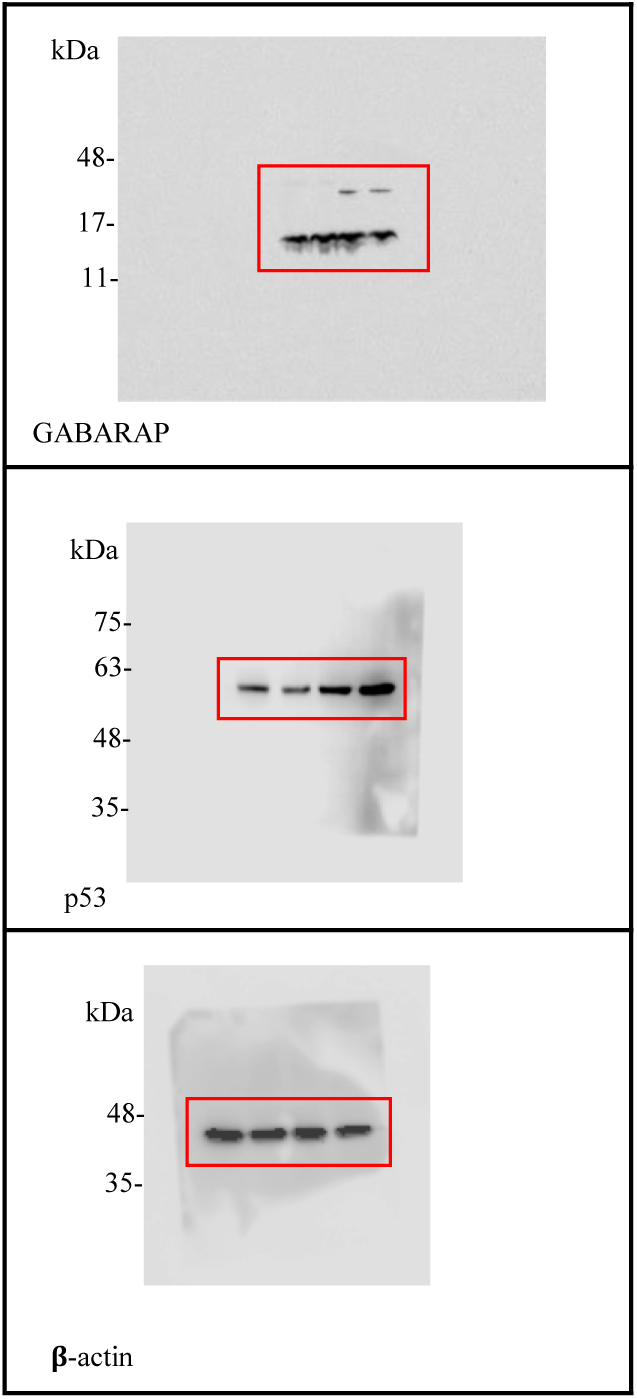

**Figure 7.**
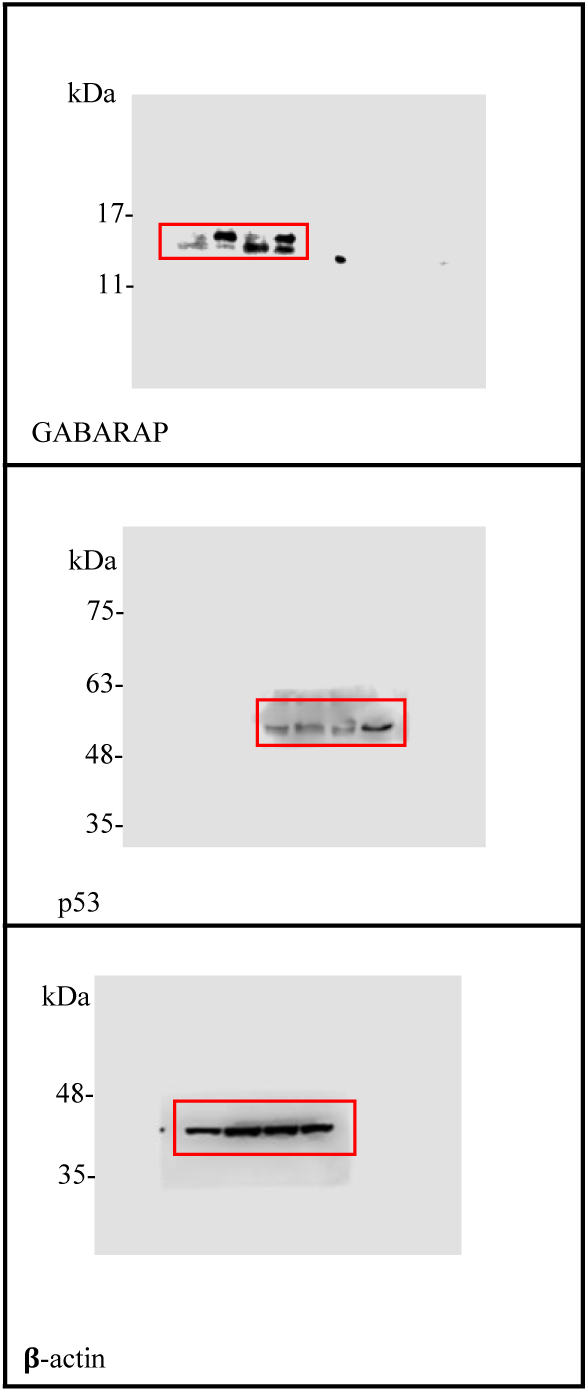

**Supp. Fig 4.**
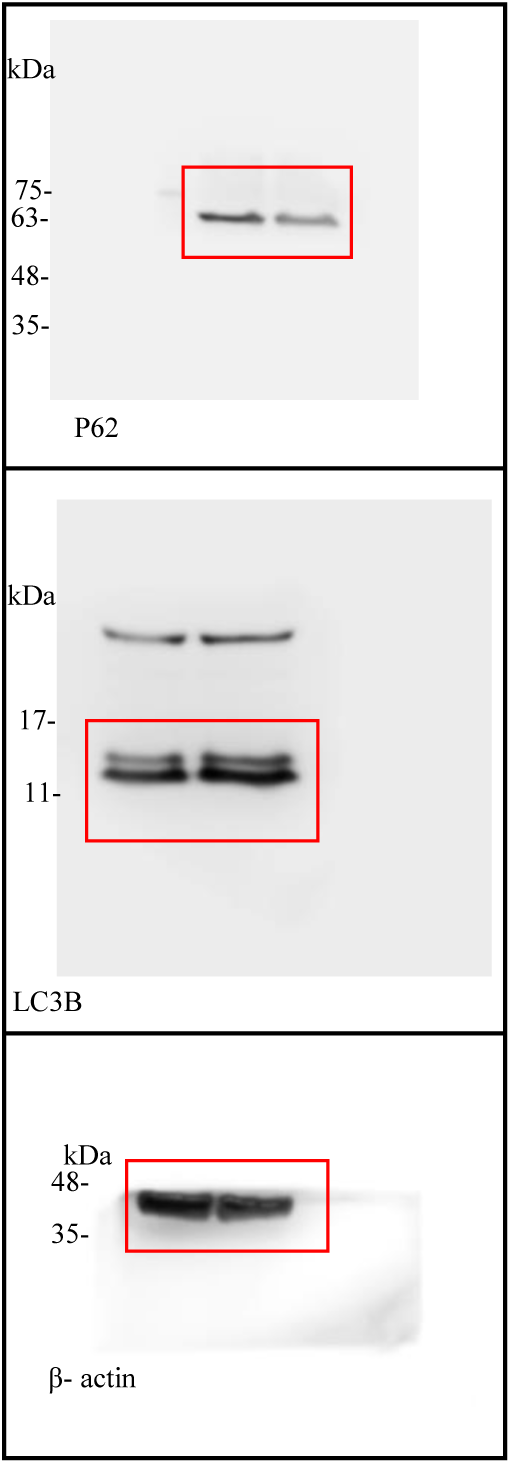

**Supp. Fig 5.**
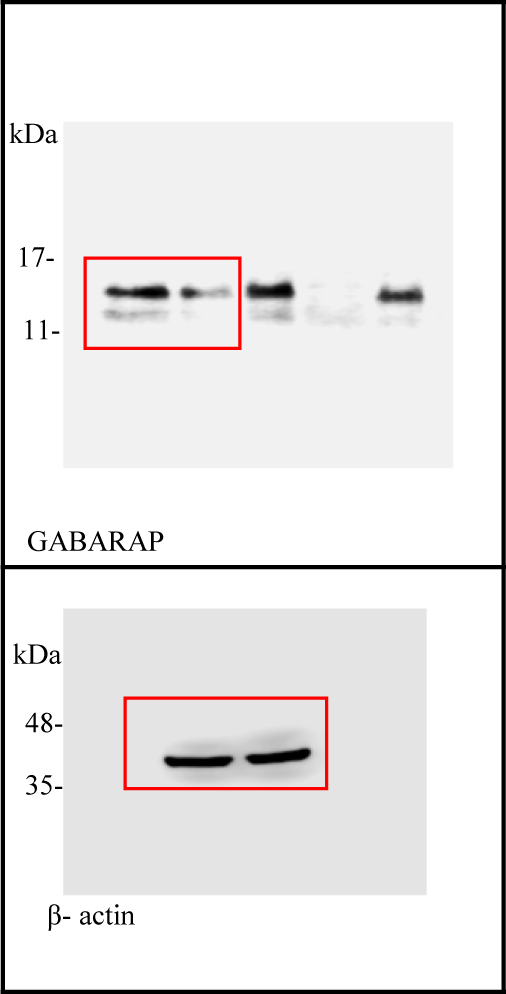

## Notes

### Competing Interest Statement

The authors have declared no competing interest.

