## Supplementary material for "Differential expression of GABARAPs in GBM renders temozolomide sensitivity in a p53-dependent manner": Table

Table 1

**GBM**

| **S.No** | **Sample code** | **Code number** | **Gender** | **Age** | **Diagnosis** |
| --- | --- | --- | --- | --- | --- |
| 1. | AIIMS-D/GBM/F/1 | 1 | F | 35 | GBM |
| 2. | AIIMS-D/GBM/F/28 | 28 | F | 58 | GBM |
| 3 | AIIMS-D/GBM/F/32 | 32 | F | 32 | GBM |
| 4 | AIIMS-D/GBM/F/34 | 34 | F | 39 | GBM |
| 5 | AIIMS-D/GBM/M/50 | 50 | M | 58 | GBM |
| 6 | AIIMS-D/GBM/M/53 | 53 | M | 30 | GBM |
| 7 | AIIMS-D/GBM/M/55 | 55 | M | 40 | GBM |
| 8 | AIIMS-D/GBM/M/62 | 62 | M | 35 | GBM |
| 9 | AIIMS-D/GBM/M/91 | 91 | M | 25 | GBM |
| 10 | AIIMS-D/GBM/F/120 | 120 | F | 18 | GBM |

**LGG**

| **S.No** | **Sample code** | **Code number** | **Gender** | **Age** | **Diagnosis** |
| --- | --- | --- | --- | --- | --- |
| 1. | AIIMS-D/LGG/F/1 | 1 | F | 37 | LGG |
| 2. | AIIMS-D/LGG/M/2 | 2 | M | 55 | LGG |
| 3 | AIIMS-D//LGG/M/3 | 3 | M | 25 | LGG |
| 4 | AIIMS-D/LGG/M/4 | 4 | M | 52 | LGG |
| 5 | AIIMS-D/LGG/M/9 | 9 | M | 25 | LGG |
| 6 | AIIMS-D/LGG/F/10 | 10 | F | 43 | LGG |
| 7 | AIIMS-D/LGG/M/13 | 13 | M | 34 | LGG |
| 8 | AIIMS-D/LGG/F/15 | 15 | F | 45 | LGG |
| 9 | AIIMS-D/LGG/F/16 | 16 | F | 27 | LGG |

**Table 2.**

| Gene | Primer | Sequence | Amplicon size |
| --- | --- | --- | --- |
| MAP1LC3A | Forward | GAACTGAGCTGCCTCTACCG | 124bp |
|  | Reverse | CCAGAGGGACAACCCTAACA |  |
| MAP1LC3B | Forward | AGCGTCTCCACACCAATCTC | 102bp |
|  | Reverse | CAATTTCATCCCGAACGTCT |  |
| MAP1LC3C | Forward | ACAAGAGGAAGTTGCTGGAATCCG | 154bp |
|  | Reverse | GATGATGCTGAGGAACTGGGTCAT |  |
| GABARAP | Forward | AGAAGAGCATCCGTTCGAGAA | 116bp |
|  | Reverse | AGAAGAGCATCCGTTCGAGAA |  |
| GABARAP L1 | Forward | CCCTCCCTTGGTTATCATCCA | 121bp |
|  | Reverse | ACTCCCACCCCACAAAATCC |  |
| GABARAPL2 | Forward | TCGAGCGAAATATCCCGACA | 179bp |
|  | Reverse | CCACAAACAGGAAGATCGCC |  |
| 18s rRNA | Forward | CGGCGACGACCCATTCGAAC | 99bp |
|  | Reverse | GAATCGAACCCTGATTCCCCGTC |  |
| P53 | Forward | AGGTTGGCTCTGACTGTACC | 140bp |
|  | Reverse | AAAGCTGTTCCGTCCCAGTA |  |

| S.No. | Si RNA | Sequence |
| --- | --- | --- |
| 1. | GABARAP siRNA (NM_007278.2) | GUUGGUCAGUUCUACUUCU  dTdT  CACCAUGAAGAAGACUUCU   dTdT  GACUUCUUUCUCUACAUUG  dTdT |
| 2. | GABARAPL1 siRNA (NM_031412.4) | GAGUGUCUAUGGGAAAUGA dTdT  UGAGGACAAUCAUGAGGAA  dTdT  UCUGGACAAGAGGAAGUAC  dTdT |
| 3. | GABARAPL2 siRNA (NM_007285.7) | GAUUGUUGACAUUGACAAA  dTdT  GUACUUGGUUCCAUCUGAU  dTdT  CAGCUUUACGAGAAGGAAA  dTdT |
| 4. | TP53 siRNA (NM_000546.6) | GCAUCUUAUCCGAGUGGAA  dTdT  CCAUCAUCACACUGGAAGA   dTdT  GAUGUUCCGAGAGCUGAAU  dTdT |

**Si RNA**
